# Running on empty: Mitochondria without DNA exhibit differential motility and connectivity

**DOI:** 10.1101/2024.08.27.609985

**Authors:** Joanna M. Chustecki, Alora Q. Schneider, Madeleine H. Faber, Alan C. Christensen

## Abstract

Plant mitochondria are in continuous motion. While providing ATP to other cellular processes, they also constantly consume ATP to move rapidly within the cell. This movement is in part related to taking up, converting and delivering metabolites and energy to and from different parts of the cell. Plant mitochondria have varying amounts of DNA even within a single cell, from none to the full mitochondrial genome. Because mitochondrial dynamics are altered in an *Arabidopsis* mutant with disrupted DNA maintenance, we hypothesised that exchanging DNA templates for repair is one of the functions of their movement and interactions. Here, we image mitochondrial DNA by two distinct methods while tracking mitochondrial position to investigate differences in the behaviour of mitochondria with and without DNA in *Arabidopsis thaliana*. In addition to staining mitochondrial DNA with SYBR Green, we have developed and implemented a fluorescent mitochondrial DNA binding protein that will also enable future understanding of mitochondrial dynamics, genome maintenance and replication. We demonstrate that mitochondria without mtDNA have altered physical behaviour and have a lower immediate connectivity to the rest of the population, further supporting a link between the physical and genetic dynamics of these complex organelles.

## Introduction

Mitochondria are highly dynamic, bioenergetic organelles. In most plant cells, mitochondrial populations exist as individual organelles, moving about the cell interacting with each other and other organelles, while remaining evenly spread across the cell (Chustecki *et al*., 2021). This movement is mostly facilitated by myosin motors upon actin filaments (Avisar et al., 2009; Romagnoli et al., 2007). This dynamic behaviour may be important for taking up, converting, and delivering metabolites and energy to different parts of the cell, and providing the opportunity to exchange mitochondrial contents. In contrast to other kingdoms, not all mitochondria in plant cells carry mitochondrial DNA (mtDNA) (Arimura *et al*., 2004; Preuten *et al*., 2010; Gualberto *et al*., 2014; Kozik *et al*., 2019). Some individuals carry the full complement of genes, or a subgenomic molecule, while others carry none. There is evidence that low levels of mtDNA in plants are autonomously regulated and are necessary to maintain RNA editing capacities and plant respiratory health (Zhang et al., 2023).

Exchange of mtDNA between mitochondria in plant cells has been demonstrated on the level of single organelles (Sheahan *et al*. 2005; Arimura, 2018), and this exchange may allow each individual to access the full complement of genes and gene products over time (Logan, 2006; Arimura, 2018; Giannakis et al., 2022). Plants also exhibit high levels of mtDNA recombination and varied physical forms of nucleoids, including linear and branched (Kozik et al., 2019). Mitochondrial interactions therefore offer another level of control for DNA maintenance, repair and recombination, and allow individuals to be interconnected without being physically joined (Johnston, 2019).

Colocalisation, fission and fusion events as well as ‘kiss-and-run’ membrane sharing dynamics have functional implications for plant mitochondria (Bauwe et al., 2010; Christensen, 2014; Hagemann et al., 2016; Midorikawa et al., 2022; Mueller & Reski, 2015; White et al., 2020). This interacting population has been demonstrated to homogenise mitochondrial contents, explicitly seen in onion epidermal cells over a few hours (Arimura et al., 2004). Therefore, the idea of a ‘discontinuous whole’, has emerged (Logan, 2006) - describing individual organelles as part of a whole interconnected population While some developmental stages and cell types have reticulated mitochondrial structure, (such as the shoot apical meristem (Segui-Simarro et al., 2008), or germinating embryos (Paszkiewicz et al., 2017)), individual mitochondria in most plant cells *transiently* and *continuously* share their contents. This link between the *physical* and *genetic* dynamics of plant mitochondria is not yet well-understood.

Disruption of the organellar DNA repair gene *MSH1* imposed a genetic challenge to the mitochondrial genome, leading to increased colocalisation times and network connectivity (Chustecki et al., 2022). This was hypothesised as a response to increased sharing due to individual mitochondria needing to access undamaged mtDNA templates for double-strand break repair (Chustecki *et al*., 2022; Rodriguez *et al*., 2022). This leads directly to the hypothesis that the mitochondrial genome is an important influence on mitochondrial dynamics and network behaviour.

To test this hypothesis we developed a system to assess whether the dynamics of mitochondria with and without DNA are different. We used a combination of *Arabidopsis thaliana* single-cell confocal microscopy, tracking of dynamic movement, and network analysis to characterise mitochondrial behaviour and correlate it to the presence or absence of DNA. To image mtDNA we use two systems, an exogenous dsDNA stain, and we also introduce a novel mtDNA binding photoconvertible fluorescent protein to Arabidopsis, providing a new tool to quantify, track and reveal mtDNA dynamics in live cells, eliminating the need for exogenous stains. We present data showing links between mitochondrial DNA, mitochondrial dynamics and social behaviour.

## Results

### Plant cells show a persistent population of mitochondria without mtDNA

It has previously been shown that individual mitochondria in plant cells do not carry the full gene complement. Some do carry the full complement, while some carry a subgenomic molecule(s) with a few genes, and some have no mtDNA at all (Arimura et al., 2004; Gualberto et al., 2014; Kozik et al., 2019; Logan, 2006; Preuten et al., 2010; Takanashi et al., 2006). We quantified the proportion of mitochondria with and without mtDNA in *whole* single cells, across two cell types, by fixing 5-day old *Arabidopsis thaliana* seedlings encoding mitochondrially-targeted mCherry (Kindly provided by Prof. Markus Schwarzländer (pBin atp-mCherry, (Candat et al., 2014))). These were stained with SYBR green, an intercalating dye specific for double stranded DNA (dsDNA), with a low affinity for RNA (Zipper *et al*., 2004; Arya *et al*., 2005; Dragan *et al*., 2012). Cells (8 hypocotyl cells and 11 leaf epidermal cells) were imaged using a confocal laser scanning microscope and segmented using IMARIS (see methods). The SYBR stain gives a signal for all dsDNA in the cell, including the nucleus and chloroplast, alongside the mitochondrial DNA signal of interest (Figure 1A.i,ii). Some organelles contain no mtDNA, while others have a range of weaker to stronger SYBR signals, all able to be detected by the segmentation thresholds implemented (Figure 1B). Only the SYBR signal that sits directly within mitochondrial volumes was quantified, avoiding other organellar DNA signals. This method revealed a persistent but varied proportion of mitochondria that do not carry genomic material, (Figure 1C, see methods), confirming previous results (Arimura et al., 2004; Zhang et al., 2023).

**Figure 1:**
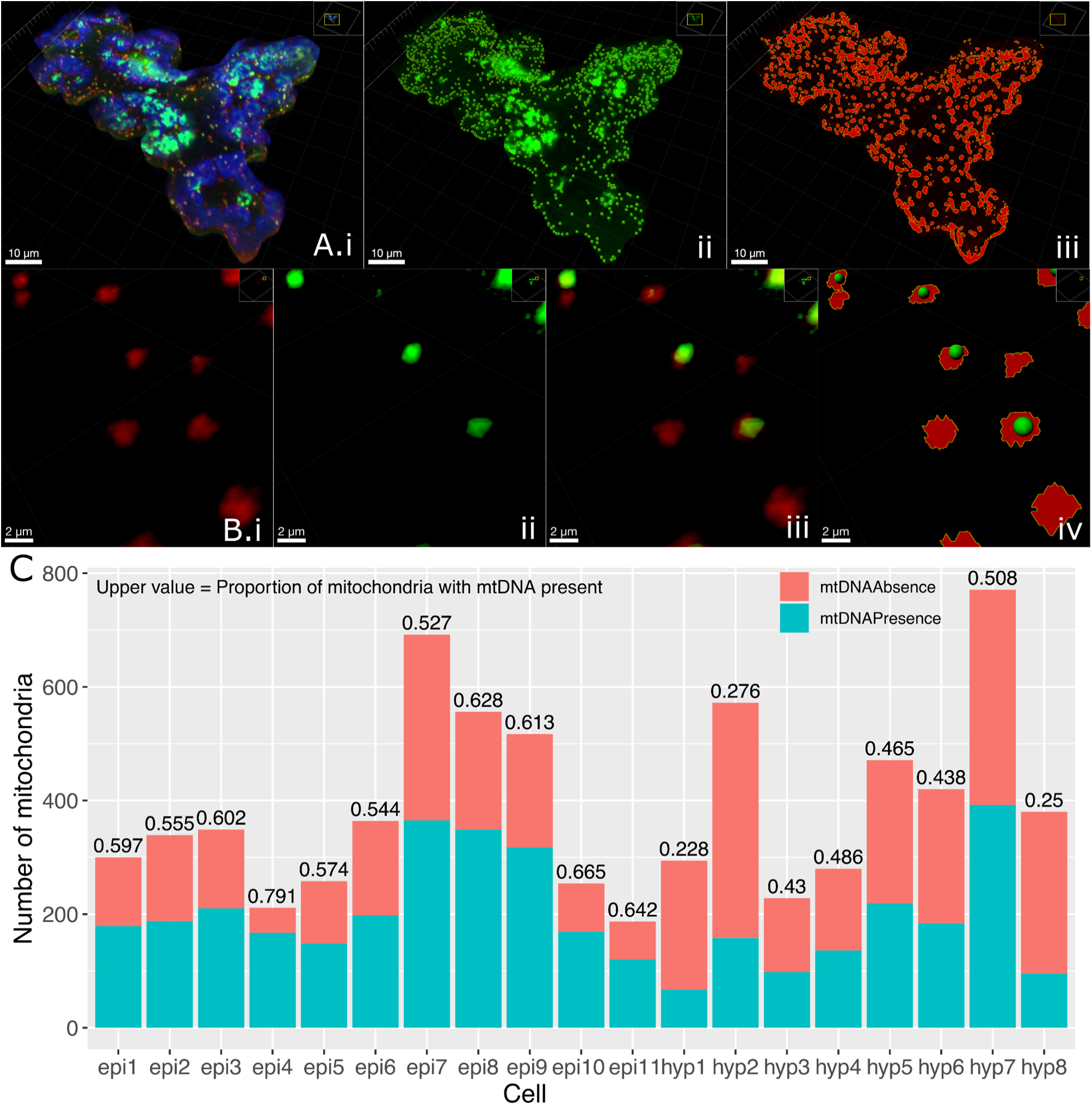
Quantification of mitochondria with and without mtDNA presence in fixed single plant cells A) (i) A single, fixed, leaf epidermal cell with mito-mCherry marker (red), SYBR green staining ds-DNA (green), and chloroplast autofluorescence (blue). (ii) SYBR channel with segmented puncta of ds-DNA signal, (iii) mito-mCherry channel with thresholded 3D rendering of mitochondria. All (A) are Z-stacks of 40 x 0.3µm slices, whole cell rendered in IMARIS. B) Magnified portion of cell shown in (A), with (i) the mito-Cherry signal, (ii) the SYBR signal, (iii) both mito-mCherry signal and SYBR signal, (iv) thresholded 3D rendering of both channels segmented using the IMARIS cell model. C) Number of mitochondria with or without mtDNA following IMARIS segmentation. Height of bars is total number of mitochondria, mtDNA absent number is in coral (upper part of bar), mtDNA present number is in blue, (lower part of bar). Value printed above bars is the proportional value of mitochondria with mtDNA present. ‘epi’= leaf epidermal cell, ‘hyp’ = hypocotyl cell.

Both the proportion and absolute number of mitochondria not carrying genomic material vary both across and within cell types. As controls for the segmentation process, a Col-0 (wild-type, with no stable fluorescent marker) hypocotyl cell treated with SYBR, and a mito-mCherry hypocotyl cell with no SYBR stain were also segmented, giving similar intensity and mitochondrial volumes, respectively (Figure S1). We also see a correlation between cell area and mitochondrial number, explaining the variation in total mitochondrial number across cells (Figure S2). This staining and segmentation offers a cell-by-cell quantification of mtDNA presence/absence, supporting previous observations of mitochondria without genomic material.

### Defining mtDNA presence or absence in moving mitochondria

To investigate the hypothesised link between the physical and genetic dynamics of plant mitochondria, we characterised the movement of individual organelles with and without mtDNA. Laser scanning confocal time lapses of 5-day old mito-mCherry hypocotyl cells stained with the dsDNA marker SYBR green were captured over ∼4 mins (Figure 2A, Video 1). Hypocotyl cells were chosen as they have been demonstrated to provide a quasi-2D field of view of the mitochondrial population, and the cellular geography allows clear tracking of individual organelles (Chustecki *et al*., 2021).

**Figure 2:**
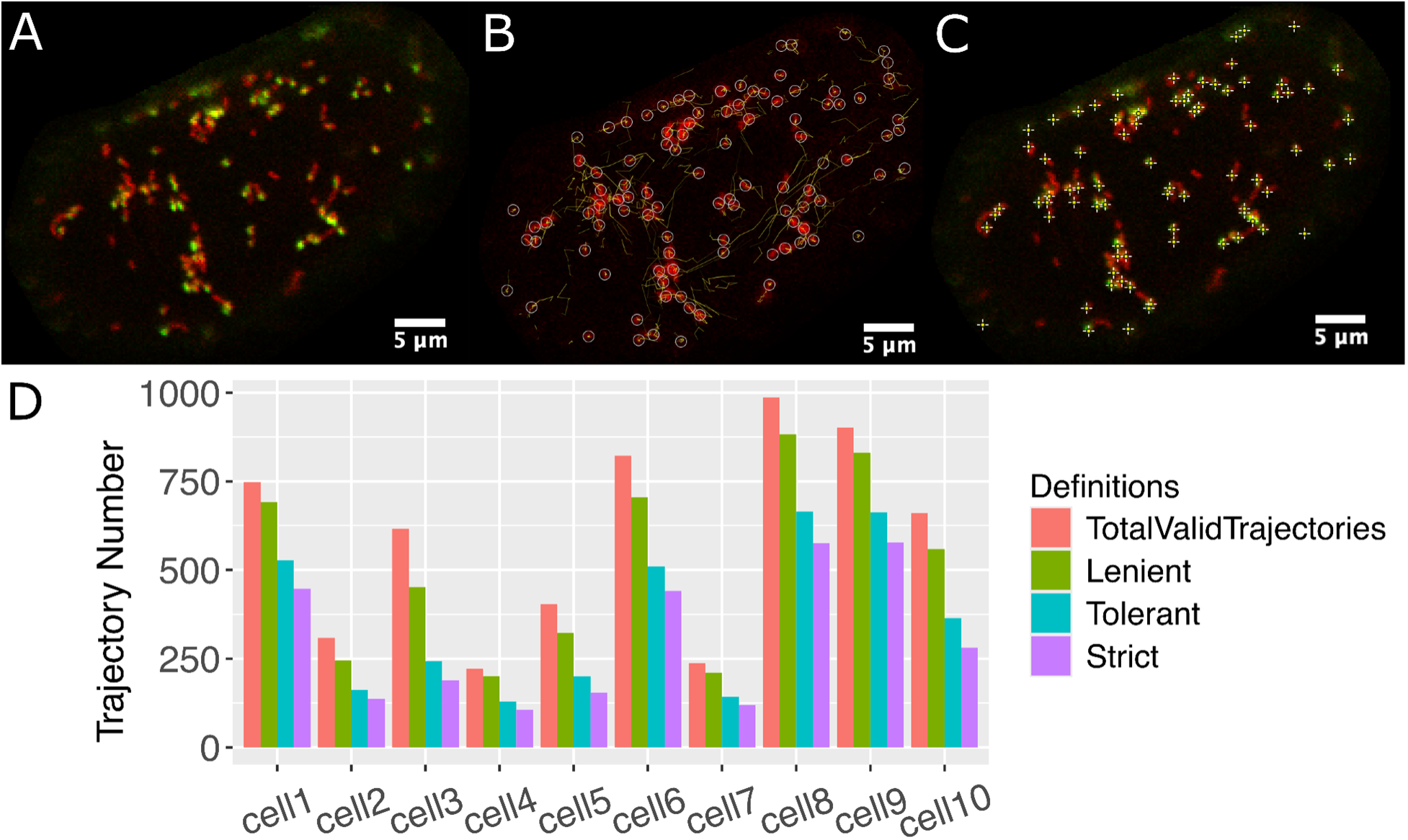
Detecting and defining mtDNA presence. A-C; Single *Arabidopsis* hypocotyl cell with mito-mCherry marker, stained with SYBR green, (A) still frame from time lapse. B) Detected mitochondria (red) using Trackmate (white circles, diameter 1.1µm) with trajectories of mitochondrial motion (yellow), limited to ten frames for clarity (5 frames either side of current position). C) SYBR green signal puncta highlighted (white crosses) upon points of intensity classified by Find Maxima function. D) Quantification of trajectory number over total, alongside three definitions of mtDNA presence, in order of strictness, across 10 single hypocotyl cells.

Tracking was performed using Trackmate (Tinevez *et al*., 2017), upon the mito-mCherry channel of the captured videos, followed by manual verification. The SYBR green channel was then used to define mtDNA presence or absence and did not influence the tracking. Trackmate quantification results can be viewed as a trajectory map over time (Figure 2B).

Two approaches were tested to quantify SYBR signal in the GFP channel. Firstly, Trackmate quantifies the signal in each dot encircling the tracked mitochondrion. From this, the Contrast statistic was used to define presence or absence of mtDNA (Tinevez et al., 2017). Contrast is calculated as the Intensity (A.U) of pixels inside the spot radius (*r*) compared to outside as defined by 2*r* by (I_in_-I_out_)/(I_in_+I_out_) (based on (Michelson, 1995)). This is a good indicator of SYBR signal for punctate mitochondria (Figure S3A). However due to the intensity considered for I_out_, two distinct SYBR signals that sit within 2*r* of each other may interfere with the contrast calculation, leading to only one being accounted for. Therefore we also use a second definition of absence/presence, inspired by the flexible parameter Prominence within mtFociCounter (Rey et al., 2023), implementing the Find Maxima function in ImageJ (Wagner, 2016), which scans each pixel intensity value, and in analogy to geographic topography, finds the maxima in relation to those surrounding it. Find Maxima distinguishes two proximal SYBR signals more effectively than the previously defined Contrast value (Figure 2C, S3A), and was carried forward as the detection method used.

The next challenge was describing the presence/absence of mtDNA across time points of individual trajectories. The complexity and speed of mitochondrial dynamics in live plant cells poses a challenge for all tracking software. Correlating SYBR signal at each time point for each mitochondrion, it was observed that organelles with no SYBR signal may occasionally have a single frame that crosses the threshold for signal. This can be seen in trajectory schemes labelled with SYBR channel features-some trajectories have a consistent SYBR signal throughout time, and others have occasional frames with a higher/lower signal (Figure S3C,D). This could be due to an unspecific SYBR signal or crossing paths with a mitochondrion with a strong SYBR signal. It could also be that those trajectories with SYBR signal may have a few frames where the signal is below the threshold (either defined by Contrast or Maxima). It is therefore necessary to implement some definition for mtDNA presence/absence, to classify this inherently noisy data.

We therefore implemented and evaluated three definitions for mtDNA ‘presence’ for trajectories - based on how many frames a mitochondrial trajectory needs to have a SYBR signal above a threshold for it to be categorised as having mtDNA. The first definition, which we will refer to as *lenient,* is that only one frame in the entire trajectory needs to be above a threshold value to define that mitochondrion as having mtDNA. The next is *tolerant*, where at least two adjacent frames must be above a threshold. The last definition is *strict*, where at least three adjacent frames must be above a threshold. This last definition necessarily excludes all trajectories that are only two frames long. Also, it should be noted, these are definitions of mtDNA presence, aiming to find a balance between accurately defining presence, while not allowing a leaky absent definition to be populated by trajectories with signal.

When quantifying the number of trajectories across each definition, the most lenient definition appears to allow too many false positives, while the strictest results in a loss of data (Figure 2D). Therefore, we settled upon the tolerant definition moving forward, however we corroborate our findings across the definitions as they are examined (see supplementary material). Paired with the Find Maxima SYBR detection method, false detection is limited, while allowing flexibility to work around any mis-tracking through the software. This allows us to confidently define and detect mtDNA per trajectory for the subsequent physical statistical analysis.

### Mitochondria without mtDNA demonstrate altered physical dynamics

Using positional data of mitochondrial movement, and the previously described *tolerant* definition of mtDNA presence/absence, physical statistics of individual mitochondria with and without mtDNA were compared (see methods). First, mean speed was calculated per trajectory, and no difference was observed between mitochondria with and without mtDNA, across ten hypocotyl cells (Figure 3A). When comparing the mean area travelled per trajectory, a significantly smaller area is travelled by mitochondria without mtDNA (Figure 3B), despite no overall difference in speed here. Area travelled is necessarily dependent both on speed, direction, and in this case, trajectory length. We see a difference in trajectory length between those with and without mtDNA, where mitochondria without mtDNA have shorter trajectory duration (Figure S4A). To investigate this further, we implemented the *most extreme* possible definition of mtDNA presence and absence: every single time point in a trajectory must pass the Maxima threshold to count as that mitochondrion having mtDNA, and every single time point in those without mtDNA must be below the Maxima threshold. This results in any trajectories where not every time point was either with or without DNA were discarded. In this case, we see similar trajectory lengths, but still a smaller area travelled by mitochondria without mtDNA, confirming that this relationship holds and rules out artefact from presence/absence definitions (Figure S4B,C). We also see this across all definitions (Figure S8)

**Figure 3:**
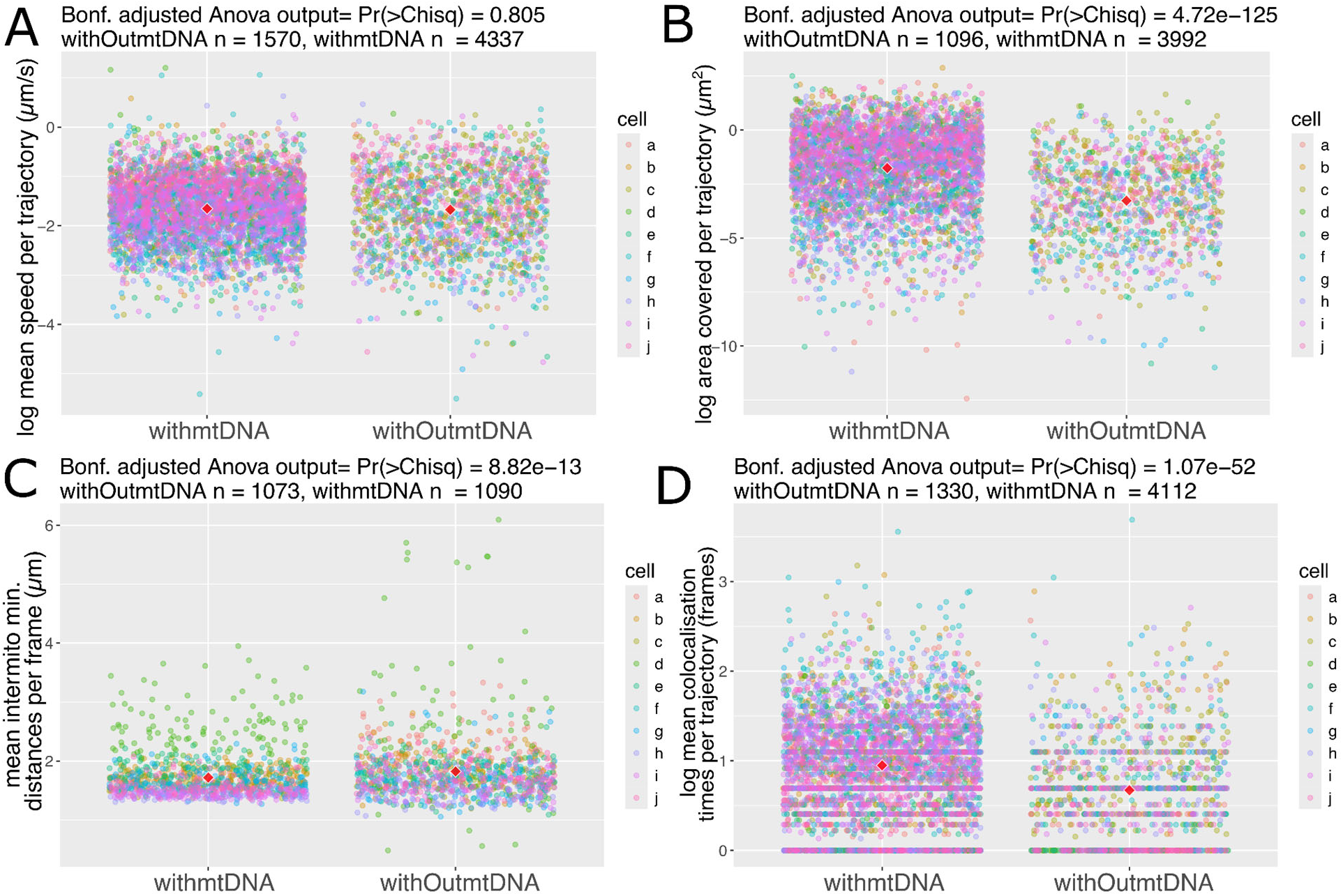
Physical statistics of mitochondrial motion compared between mitochondria defined as with/without mtDNA, across n= 10 SYBR stained hypocotyl cells over 240 seconds of frame time A) Mean speed calculated per trajectory (µm/s) B) Convex hull, the area inside a polygon of the furthest points in a trajectory (µm^2^) C) Minimum Euclidean distance between a single mitochondrion and all others, averaged per frame for with/without mtDNA (µm) D) Number of frames individual mitochondria are colocalised to any other. P-values are the outcome of a blocking factor Anova (cells as block), adjusted across all statistics tested using the Bonferroni method. All plots use the *tolerant* definition of mtDNA presence/absence. Red diamonds represent the mean of all points.

Returning to our *tolerant* definition of mtDNA presence/absence, when comparing intermitochondrial positioning, we see differences between mitochondria with and without mtDNA. Intermitochondrial distance is the minimum Euclidean distance between each mitochondrion in each trajectory, averaged per frame. Mitochondria without mtDNA show an increased intermitochondrial distance, meaning they are on average further away from every other mitochondrion in the system than those with mtDNA (Figure 3C). Following this, to test the propensity for sharing between mitochondria, colocalisation time was calculated as the number of frames mitochondria had been within 1.6µm distance of each other (as in (Chustecki *et al*., 2021)). There is a shorter colocalization time between those without mtDNA and any other mitochondrion it comes into close contact with, than those that do have mtDNA (Figure 3D).

Examination of the same physical statistics across the two foci definitions (Contrast/Maxima) and the three trajectory definitions (Tolerant/Lenient/Strict) reveals consistent conclusions to the Maxima and *tolerant* used to define mtDNA presence here (Figure S8). Overall, a difference in movement between those with mtDNA and without mtDNA is observed, across whole populations of mitochondria in single cells.

### Mitochondria without mtDNA are less well connected to the rest of the population

To study mitochondrial connectivity across whole populations of individual organelles, a novel method borrowing from graph theory, analogous to social networks, has been developed (Chustecki *et al*., 2021). Defining individual mitochondria as nodes (here represented as circles), and close connectivity (within 1.6µm distance) as a connecting edge (represented as sticks), we can successfully build historical networks describing and quantifying the encounters of individual organelles over time lapse videos of motion in single cells. Onto this, we project for the first time, functional data - the presence or absence of mtDNA. We define each trajectory as with/without mtDNA (using the Maxima, and *tolerant* definitions) and build an interaction network over time, from all mitochondria and all connections (Figure 4A).

**Figure 4:**
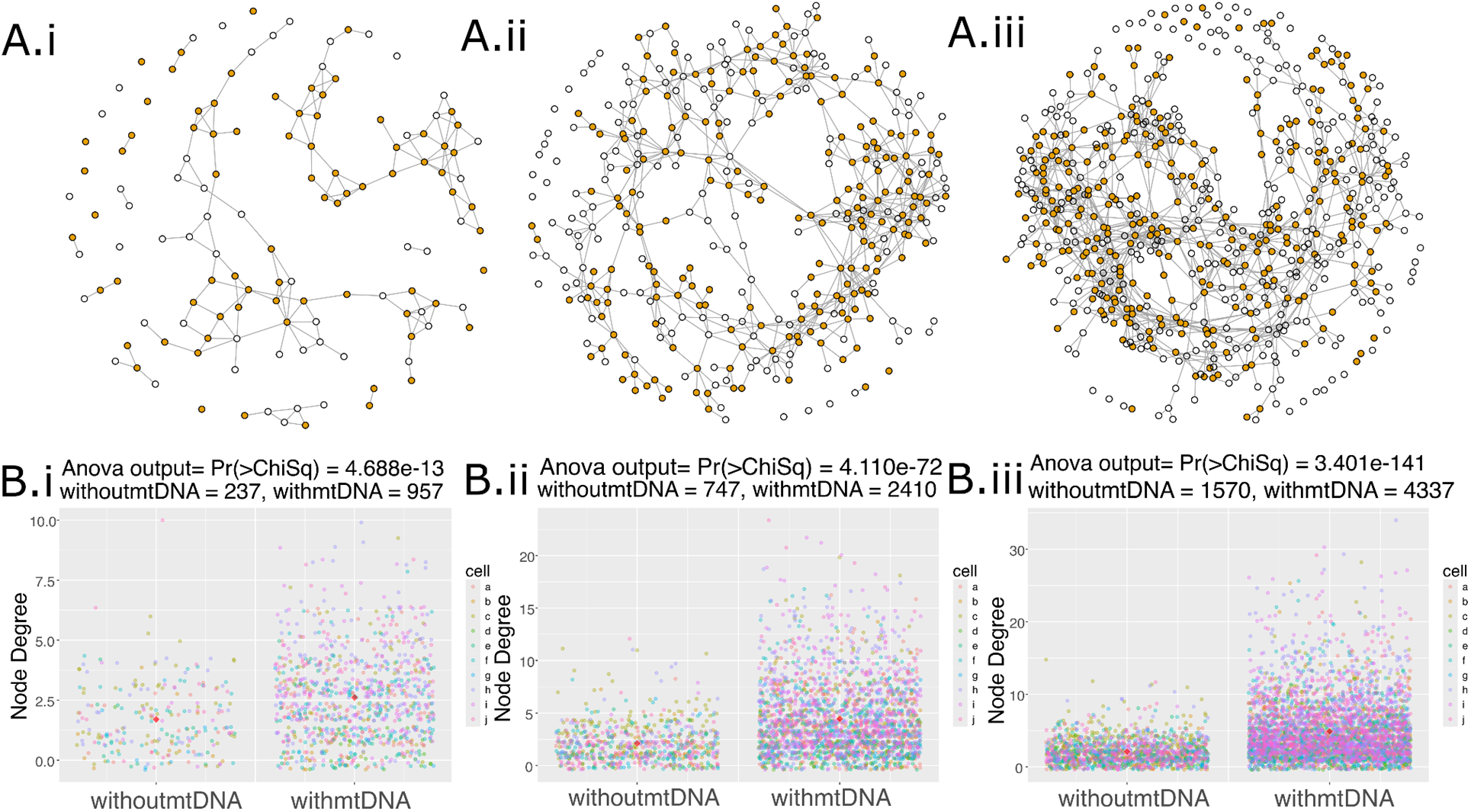
Mitochondria without mtDNA demonstrate lower immediate connectivity. A) Representative network of all mitochondrial encounters built up over time in SYBR stained hypocotyl cells, shown at frame 10 (representing 22.1sec of video time) (i), 50 (ii), 109 (iii), colour coded by mtDNA presence (orange) or absence (white), with edges representing close encounters between individual organelles. B) Node degree with mean (red diamonds) for nodes with and without mtDNA over the same frames, and across all 10 hypocotyl cells (a-j). P-values are the outcome of a two-way blocking factors anova (cells as block). Red diamonds represent the mean of all points.

Graph theory and social network analyses use a parameter called “degree” to describe the connectivity between individuals in a network. A low degree means that during the time examined an individual encountered a very small number of other individuals in the network. For mitochondria, this would mean they travel around the cell without coming into close proximity with many other mitochondria. In contrast, a high degree means that a mitochondrion was in close proximity to many others over time. In order to compare connectivity between mitochondria with and without mtDNA we focused on this node-specific value of degree, which is a measure of how many other mitochondria a single mitochondrion met over the time we tracked it within the cell, or in network graphical terms, how many sticks are connected to the circles. Strikingly, the node values for mitochondria with DNA reach much higher local connectivity levels than those without mtDNA (Figure 4). This means that mitochondria with mtDNA have more immediate neighbours (of either type) than those without.

By the final frame of all videos, we see mitochondria without mtDNA only reaching a maximum degree value of 15, while those with mtDNA reach a maximum degree value of 34 (Figure 4B.iii). Examination of the degree parameter across the two foci definitions (Contrast/Maxima) and three trajectory definitions (Tolerant/Lenient/Strict) reveals the same significant relationship of higher degree values for those with mtDNA, regardless of the definition used (Figure S9). The combination of these 12 different definitions and analysis strongly suggest that our choices of how to define a mitochondrial image and how to decide whether it has mtDNA in it or not do not change the interpretation, and the results are robust to possible artefacts of the definitions used during image analysis.

Examination of the network illustrations (Figure 4Ai-iii) suggests that those without mtDNA are enriched in those with degree 0 (i.e no close encounters with other mitochondria) and by those with degree 1 (i.e at the periphery of the network). This is confirmed by quantifying the number of nodes with a certain degree value in proportion to the number of nodes of each status at representative time points across all cells analysed (Figure 5). For those nodes of degree 0 or 1, we see a higher proportion of nodes without mtDNA than those with (Figure 5A,B). When compared to those of degree 3, i.e with three immediate neighbours (Figure 5D), which is well represented in both those with and without mtDNA (Figure 4B), we see no particular mtDNA status as prominent over frames or cells, in contrast to the representation of those without mtDNA in the lowest degree nodes of 0, 1 or 2 (Figure 5A,B). At higher degree values, above 3 (Figure 5E-I), we see a higher proportion of nodes with mtDNA compared with every other node of the same mtDNA status. Above degree 8 comparison between nodes with and without mtDNA becomes difficult due to a lack of nodes without mtDNA and with high degree (Figure 4B). When this analysis is extended to all frames of all cells, the same pattern is observed (Figure S7). Thus, mitochondria without mtDNA do not have as many immediate neighbours as those with mtDNA, and are more likely to be nodes with only zero or one immediate neighbour(s), while those with higher degree values are more likely to be mitochondria with mtDNA.

**Figure 5:**
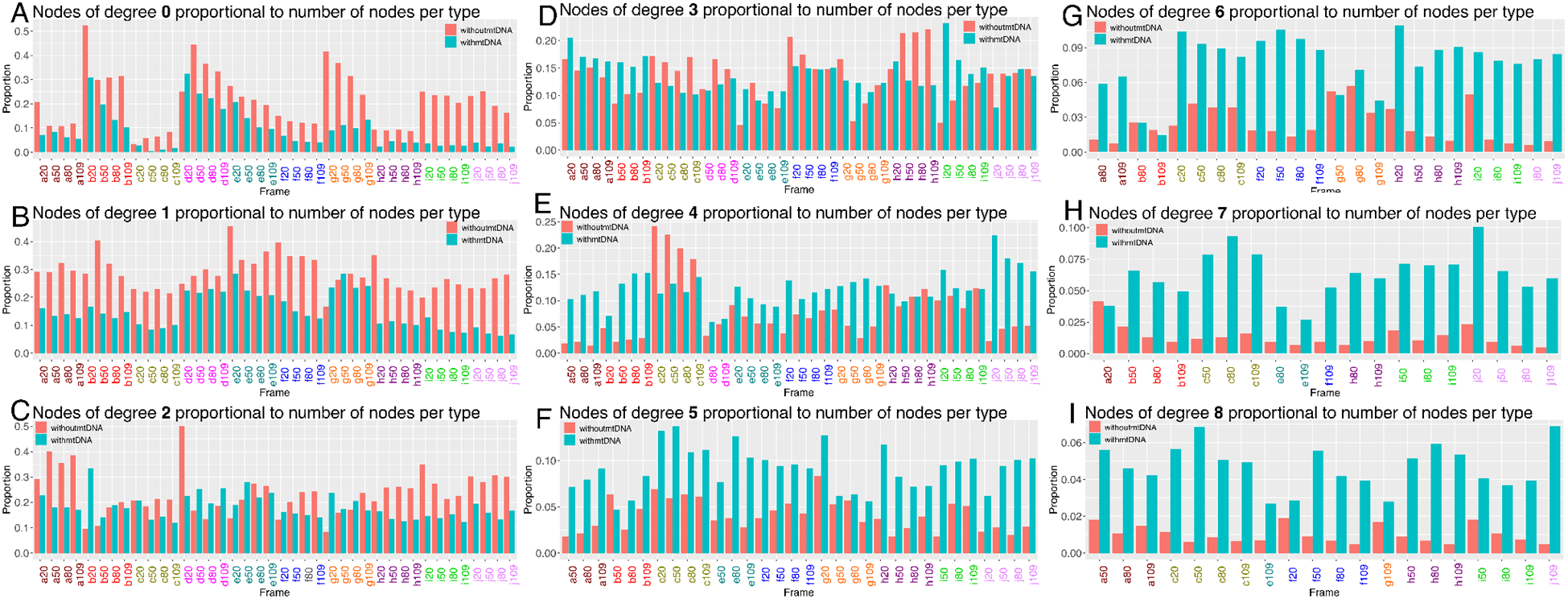
Number of nodes with degree value 0-8 (A-I), proportional to the number of nodes of equal mtDNA status, plotted over n = 10 SYBR stained hypocotyl cells, colour coded on the x-axis, and four networks at specified frames 20, 50, 80 and 109.

### Mt-HI-NESS nucleoid binding provides improved visualisation of mtDNA nucleoids in plants

Until now, mtDNA imaging in plants has relied on exogenous stains that face the impermeability barrier of the plant cell wall or fluorescent proteins that are target-specific and do not bind mtDNA ubiquitously. Live rates of mtDNA exchange across mitochondrial populations in plants are also yet to be established. Therefore, a photoconvertible ubiquitous mtDNA marker was sought. We modified the mt-HI-NESS (Mitochondrial HNS-based Indicator for Nucleic Acid Stainings) construct developed in human cells for stable transformation and ubiquitous mitochondrial targeting in plants (Deng et al., 2023). This construct relies on the H-NS bacterial nucleoid binding protein attached via a linker to the photoconvertible Kaede protein. We codon-optimised for Arabidopsis and chose two plant-specific mitochondrial targeting sequences to test, IVD and AOX (see methods). As a further test, we also used the original human Cox8 presequence construct to see whether it would function in a plant system.

Initial tests were carried out using the *Nicotiana benthamiana/Agrobacterium tumefaciens* transient transfection system (see methods), with IVD and AOX mt-HI-NESS specifically localising *within* the individual mitochondria of epidermal cells (Figure S5D,E), using a red fluorescent IVD-FP611 construct as a mitochondrial marker. Compared to the same tissue using IVD-FP611 and stained with SYBR green, we see a similar nucleoid signal, showing that mt-HI-NESS is ubiquitously staining mtDNA nucleoids (Figure S5B,D,E). Using Cox8-mt-HI-NESS, we observed non-specific chloroplast and nuclear DNA targeting (Figure S5C), presumably because the mitochondrial targeting sequence is not plant-specific.

Following successful testing in *N. benthamiana,* the AOX/IVD-mt-HI-NESS constructs were moved into Arabidopsis plants. Floral dip of mito-mCherry and Col-0 plants was performed (see methods), and positive transformants were isolated. Each combination demonstrated mitochondrial nucleoid specific binding, and here we show mito-mCherry with AOX-mt-HI-NESS (Figure 6A, B) and Col-0 with IVD-mt-HI-NESS (Figure 6D). This binding was identical to the same tissue stained with SYBR green (Figure 6C), although the exogenously applied SYBR is also visible around the outside of the cell. We did not find any differences in localisation or expression between IVD and AOX mt-HINESS (Video 2,3).

**Figure 6:**
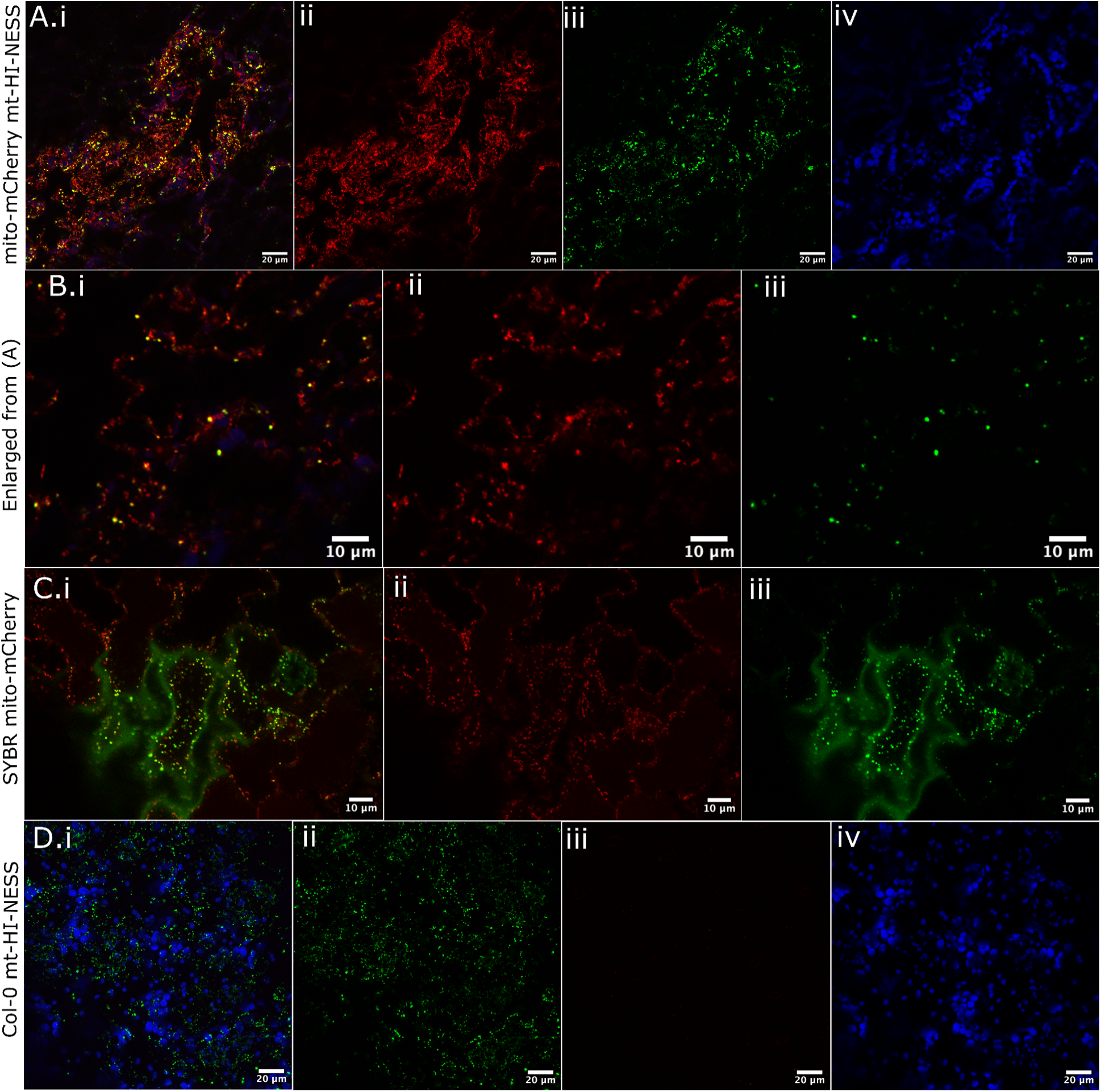
Mt-HI-NESS stable Arabidopsis transformants in mito-mCherry or Col-0 backgrounds demonstrate mitochondrial nucleoid localisation. A) Merged (i) maximum intensity projection (10 frames, 0.5µm step) with mito-mCherry (ii, red), AOX-mt-HI-NESS non-photoconverted Kaede (iii, green) and chlorophyll autofluorescence (iv, blue). B) Enlarged view of a single z-slice of same image, tiles as before. C) Single frame of mito-mCherry (red, ii), stained with SYBR Green (green, iii). D) Merged (i) Maximum intensity projection (57 frames, 0.8µm step) of Col-0 IVD-mt-HI-NESS (ii, green) with no mito marker (iii, red), and chlorophyll autofluorescence (iv, blue). All images were taken from abaxial epidermal leaf cells of mature Arabidopsis (∼2-3 weeks).

We tested the photoconvertible properties of the Kaede protein under UV laser (405nm), and demonstrated complete green-to-red switching of each mitochondrial nucleoid in a specified region of interest (Video 4).

To investigate if the IVD/AOX-mt-HI-NESS lines impact mitochondrial function through their nucleoid binding, a series of physiological experiments were conducted. The rosette areas of transformed Arabidopsis did not differ from WT (Figure S6A,B). We initially saw a difference in root length, a key phenotype for mitochondrial function, when testing Col-0 against mito-mCherry AOX or IVD-mt-HI-NESS (Figure S6C.i,ii). However, subsequent testing of the mito-mCherry line (into which these constructs had been transformed) against Col-0 revealed the same shorter root phenotype (Figure S6C.iii,E). Therefore, the difference in root length is due to the mito-mCherry background, and not the mt-HI-NESS nucleoid binding protein. This was confirmed by the root lengths in Col-0 background mt-HI-NESS lines and Col-0 showing no significant difference (Figure S6C.iv,F). Next, oxygen consumption assays as an indicator of mitochondrial respiration were performed on leaf discs in the dark on the three genotypes, and no difference in oxygen consumption rate was observed between them (Figure S6D). Finally, to determine whether the mtDNA binding ability of mt-HI-NESS disrupted expression of key genes, we used immunoblotting to assay the mitochondrial proteins Cox2 and AtpB using nuclear (Histone H3) and chloroplast (PsbA) proteins alongside as controls. We saw no differences between loading controls and signal across genotypes tested (Figure S6G). We also note that during the development of the mt-HI-NESS lines in human cells, no transcription alterations were observed (Deng et al., 2023).

### Mitochondria without mtDNA demonstrate altered motility and connectivity in a stable nucleoid marker line

Upon the validation of this new stable fluorescent mitochondrial nucleoid marker line, we performed live imaging, tracking and differential motility analysis on 11 mito-mCherry AOX-mt-HI-NESS hypocotyl cells (Figure 7). This system offers a precise and ubiquitous marking of mtDNA in plant cells, and together with the mito-mCherry marker, allows visualisation of mitochondria with and without mtDNA in a cell with very little background signal, as was often seen in the exogenous SYBR stained cells (Figure 2A, 7A). The clear signal provided in the stable genetic line is amenable to precise thresholding using the Find Maxima function used to define mtDNA puncta in the time-lapse images (Figure 7A.iii).

**Figure 7:**
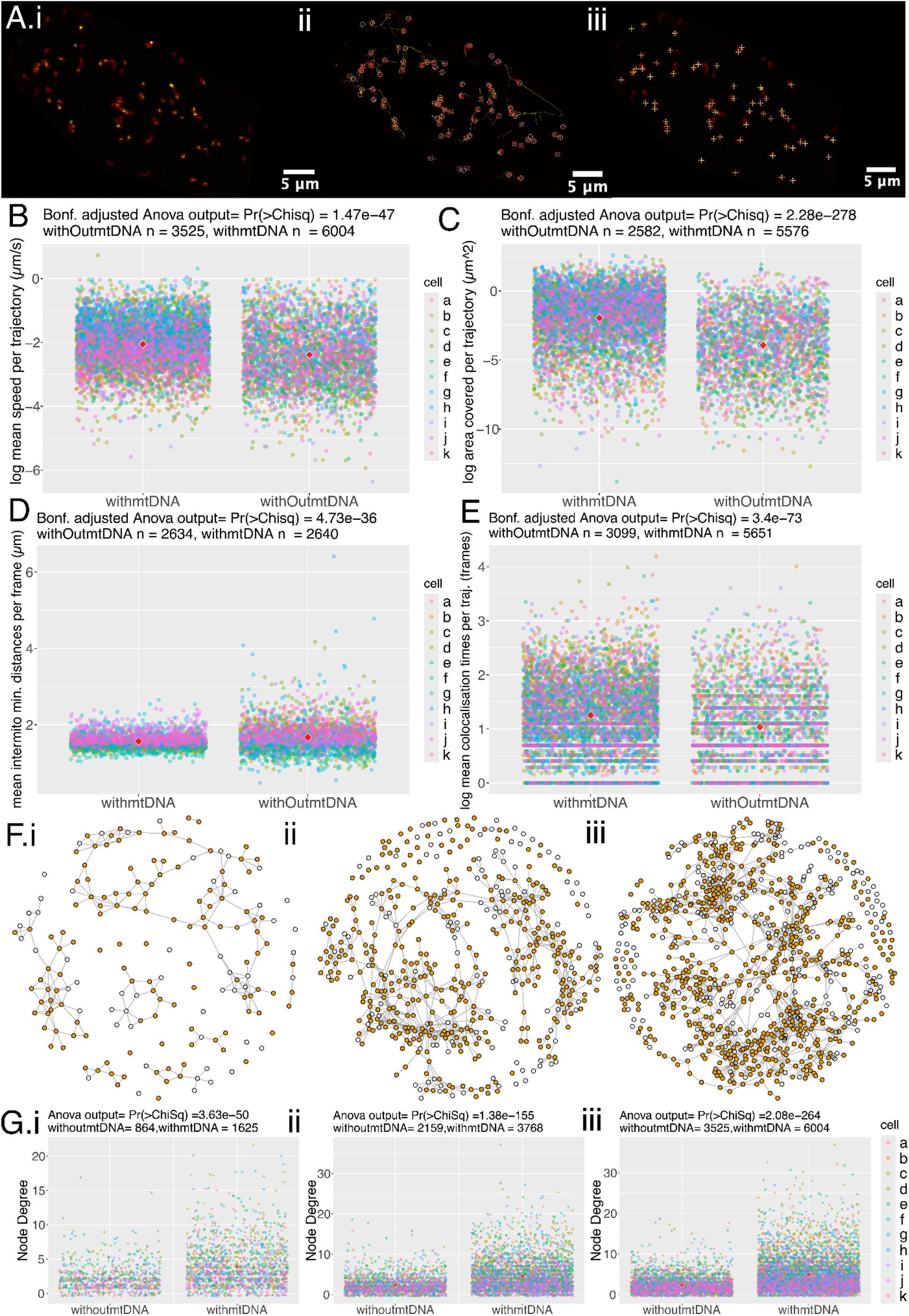
Mitochondria in mt-HI-NESS stable nucleoid marker lines also demonstrate altered physical dynamics and lower immediate connectivity in mitochondria without mtDNA compared to those with mtNDA. A) i-iii; Single Arabidopsis hypocotyl cell with mito-mCherry marker, and AOX-mt-HI-NESS mtDNA fluorescent binding protein, still frame from time lapse. ii) Detected mitochondria (red) using Trackmate (white circles, diameter 0.8µm) with trajectories of mitochondrial motion (yellow), limited to ten frames for clarity (5 frames either side of current position). iii) SYBR green signal puncta highlighted (white crosses) upon points of intensity classified by Find Maxima function. B) Mean speed calculated per trajectory (µm/s) C) Convex hull, the area inside a polygon of the furthest points in a trajectory (µm^2^) D) Minimum Euclidean distance between a single mitochondrion and all others, averaged per frame for with/without mtDNA (µm). E) Number of frames individual mitochondria are colocalised to any other. F) Representative network of all mitochondrial encounters built up over time in mt-HI-NESS hypocotyl cells, shown at frame 50 (representing 126 sec of video time) (i), 150 (ii), 240 (iii), colour coded by mtDNA presence (orange) or absence (white), with edges representing close encounters between individual organelles. G) Node degree with mean for nodes with and without mtDNA over the same frames, and across all 11 hypocotyl cells (a-k). All plots use the *tolerant* definition of mtDNA presence/absence. P-values for B-E are the outcome of a blocking factor Anova (cells as block), adjusted across all statistics tested using the Bonferroni method. P-values for G i-iii are the outcome of a two-way blocking factors anova (cells as block). Red diamonds in plots represent the mean of all points.

Upon analysis of the physical dynamics of mitochondrial motion between mitochondria with and without mtDNA in this new system, the significant relationships remain the same as seen in the SYBR system. Mitochondria without mtDNA travel around a significantly smaller area in the cell on average than those with mtDNA (Figure 7C). Those without mtDNA also demonstrate an increased intermitochondrial distance between each individual without mtDNA and the rest of the mitochondrial population (Figure 7D). This in turn is matched by a decreased colocalisation time for mitochondria without mtDNA, in that each mitochondrion without mtDNA does not spend as many frames associated with other mitochondria as those with mtDNA do (Figure 7E). For each of these conclusions, we tested the relationship across the two definitions for mtDNA foci, Contrast and Maxima, and the three definitions for trajectories (tolerant/lenient/strict) with and without mtDNA (Figure S10). These relationships are consistent across all definitions tested, with the exception of tolerant and strict definition for the contrast-detected foci in the intermitochondrial mean data, although the variation seen in those mitochondria without mtDNA remains consistent (Figure S10B.iii, C.iii) and matches that seen in the SYBR system (Figure S8B.iii, C.iii).

Interestingly, we see that mitochondria without mtDNA have a reduced speed in this mt-HI-NESS system (Figure 7B), and this conclusion is consistent across the definitions of foci and trajectories (Figure S10.A-F.i). This difference was not observed in the majority of the comparisons across definitions in the SYBR system, although some did indeed pass the threshold of significance (0.001) (Figure 3A, S9.A-F.i). We do not see the opposite conclusion anywhere, of mitochondria without mtDNA having an increased speed. The slower speeds in mitochondria without mtDNA may also be reflected in the smaller area travelled by mitochondria without mtDNA (Figure 7C). Using the mt-HI-NESS system we see significantly slower mitochondrial motion in those without mtDNA.

Upon network analysis of local connectivity between organelles in the population, mitochondria marked as having nucleoids by mt-HI-NESS again have much higher connectivity as measured by degree values than those without mtDNA (Figure 7F,G). We also see that mitochondria without mtDNA are more likely in nodes with lower degree values (degree 0,1,2) than higher degree values (3-8) (Figure S12). This confirms the relationship seen in the SYBR treated cells, and is seen consistently across all definitions of foci and trajectories (Figure S11).

Overall, the new mt-HI-NESS system provides a more specific marker of mtDNA with less background, enabling a clear-cut analysis of presence or absence of mitochondria nucleoids per organelle. Upon analysis of physical and social dynamics in the SYBR and mt-HI-NESS systems the same conclusions of differential motility and connectivity can be drawn, with the addition of a significant difference in speed.

## Discussion

Mitochondria in plant cells are highly dynamic and have many varied functions, including the generation of ATP via the electron transport chain. The most energetically and physically central proteins to this process are encoded in the mitochondrial genome, which is organised into highly variable and recombinationally active nucleoids distributed throughout the mitochondrial population. We demonstrate here a difference in both physical and network (or ‘social’) dynamics between those that carry these nucleoids, and those that do not.

Across the two different systems used, mitochondria without mtDNA have an increased intermitochondrial distance, that is, they are further apart from every other mitochondrion on average than those with mtDNA. In addition, mitochondria without mtDNA travel a smaller area around the cell than those with mtDNA. We see only two instances of significant differences in speed across definitions in the SYBR system, but we see significantly slower mitochondria without mtDNA in the mt-HI-NESS system, and across all definitions. In mitochondria with compromised mtDNA repair machinery, speed did not differ significantly to wild-type (Chustecki et al., 2022). Mitochondrial motion in plant cells is driven by myosin motors on actin filaments (Avisar et al., 2008, 2009; Sparkes et al., 2008; Zheng et al., 2010), and by cytoplasmic streaming which is a consequence of organelle motion (Shimmen and Yokota, 2004; Peremyslov *et al*., 2015; Tominaga and Ito, 2015). It may be that functional differences in mitochondria, including the presence of mtDNA do not alter this cellular machinery, although there is a core of evidence in humans and yeast that supports a trans-membrane communication between nucleoids and the cytoskeleton (Boldogh & Pon, 2006; Hoppins et al., 2007; Iborra et al., 2003; Kucej & Butow, 2007). There is no direct evidence of this yet in plants, however the influence of actin filaments in inheritance of mtDNA may instead be internal to mitochondria, as shown in mung bean (Lo et al., 2011) However, as we also see a reduced area travelled in those that do not have mtDNA, it may be that mitochondrial motion is in part arrested by colocalisation with other organelles.

Therefore, other external partners besides myosin could be involved. In animal mitochondria there is evidence of a link between ATAD3 proteins, which we now know as involved in plant mitochondrial nucleoid organisation, and the function of protein complexes at the mitochondrion-ER interface (Baudier, 2018; Kim et al., 2021). There is increasing evidence of the role of the ER in regulating mitochondrial fission and fusion (Mueller & Reski, 2015; White et al., 2020), and future research will be necessary to determine if there is indeed a link between mitochondrial nucleoids, organelle dynamics and other cellular players in these interactions.

‘Empty’ mitochondria do not have as many close connections (have a lower degree) as those with mtDNA, across any frame time, and are more likely to have 0 or 1 immediate local neighbours than those with mtDNA. Encounters between mitochondria are necessarily driven by their physical dynamics, and this behaviour correlates with the physical measurements we see for those without mtDNA.

The connection between absence of mtDNA and behaviour is difficult to disentangle. Our initial hypothesis was that empty mitochondria would make more connections with other mitochondria, in order to fuse and acquire DNA. This might happen, but we were unable to observe it, possibly because the time frame of <10 minutes is too short to see such events. Longer time frames were not done to avoid issues of photobleaching the fluorescent markers, and the plant cells becoming hypoxic under the cover slip. Why then, do empty mitochondria make fewer connections with other mitochondria? They may still be looking for partners for fusion, but mitochondria with DNA may vary in their receptivity to fusion with an empty partner, perhaps depending on how much of the genome they carry.

An additional question is why there is a persistent subpopulation of empty mitochondria. One hypothesis for this may be that those without mtDNA are targeted for degradation, and as a consequence, are also excluded from the connectivity enjoyed by the rest of the individuals in the network. However, a population of ‘empty’ mitochondria is present across all cells and both cell types tested here, as well as in all non-reproductive plant tissues where mtDNA has been imaged (Arimura et al., 2004; Kim et al., 2021; Sheahan et al., 2005; Takanashi et al., 2006). How then, is this population maintained, whether active or accidental? Upon full fusion of mitochondria, mtDNA molecules are evenly spaced over the network in plant protoplasts prior to cell division (Sheahan 2005). Thus it remains curious that this population of empty mitochondria persists. However, over the course of 2 hours in non-dividing onion epidermal cells, mitochondrial contents can be completely homogenised through fusion, and during fission events mitochondria without nucleoids can be generated (Arimura et al., 2004).

There is evidence from animals and yeast that mitochondria targeted for autophagy cease fusing to other mitochondria a few hours before degradation (Ashrafi & Schwarz, 2013; Twig et al., 2008). Interestingly, overexpression of mtSSB to force high mtDNA copy number to levels similar to those seen in animal cells is detrimental to the plant, and there is evidence that plants must retain this low copy number to maintain proper RNA editing (Zhang et al., 2023). Therefore, it seems that mtDNA-less mitochondria are continuously maintained, and non necessarily targeted for degradation.. The gain and loss of mitochondrial DNA within the population of mitochondria in a single cell is clearly a very active and dynamic process, worth further study. We know little about mtDNA homogenisation and exchange in non-reproductive tissues, and tools such as the photoconvertible mt-HI-NESS probe developed here will help us investigate further.

In non-reproductive cells, the maintenance of organellar DNA is reduced or eliminated, resulting in overall loss of DNA in older cells (Kumar et al., 2014; Oldenburg & Bendich, 2015). To avoid this potential problem, our experiments used very young plants, including leaf epidermal cells that will form much larger rosette leaves and will be metabolically active for another two months. Furthermore the previously described effect of an *msh1* mutant on mitochondrial dynamics shows that the plant is still maintaining organellar DNA with nuclear-encoded repair proteins in these cell types and stages (Chustecki et al., 2022). For these reasons, we do not think there is organellar DNA abandonment or decay in these experiments.

Defining whether a given mitochondrion has DNA or not given the complex dynamics of plant mitochondria over time is challenging. Tracking multiple shapes over time in video data using more than one channel as input is also a non-trivial problem in practice (Hänselmann & Herten, 2017; Qiang et al., 2017; Spilger et al., 2022). Many tools designed to quantify mtDNA have been developed in human or yeast cells, where mitochondria form reticulated lengths, or in only single frame images which are not suitable for this analysis (Kukat et al., 2011; Rey et al., 2023). We took several approaches to define ‘with and without DNA’ and chose the strategy resulting in the least data loss while avoiding false positives, while demonstrating that these definitions do not alter the main conclusions drawn from this analysis (Figures S8-11). It is also possible to define single time points in each video as present/absent, negating the trajectories definitions of lenient/tolerant/strict, and although we did not include this analysis here, we could not find evidence of DNA exchange from a mitochondrion with mtDNA to one without. This may be because the imaging time (<10 minutes) was not long enough to locate these events. It may also be that ‘empty’ mitochondria do not often receive nucleoids from their neighbours.

It has been demonstrated that fusion requires a potential across the inner membrane in plant protoplasts (Sheahan 2005), and that mitochondrial fusion apparatus adapts in response to changes in membrane potential, at least in mammalian cells (Herlan et al., 2004; Twig et al., 2008). Therefore, it could be the case that those without mtDNA do not have the machinery to maintain membrane potential, therefore reducing their ability to fuse. This raises the question, how do mitochondria without mtDNA keep their protein apparatus, subject to constant degradation, healthy enough to participate in this process of fusion and fission? The answer is most likely the exchange of other gene products, or whole membrane protein complexes and RNAs, during transient close proximity and full fission and fusion events, rather than the requirement to carry mtDNA at all times (Giannakis, 2022). Analysis of other functional molecules and their exchange or homogenization over these timescales would allow investigation of this, as well as analysis of any correlation between membrane potential and mtDNA presence/absence, or the establishment of functional subpopulations of these organelles (Kuznetsov *et al*., 2006; Logan, 2006; Kuznetsov *et al*. 2009; Fuchs *et al*., 2016; Willingham *et al*., 2021).

Most plant cells do not readily take up exogenously applied stains and dyes, requiring long stain times and high dye concentrations to be successful. Exogenous DNA stains are nonspecific, staining chloroplast and nuclear DNA, and can intercalate DNA molecules, impacting replication (Prole, 2020; Deng *et al*., 2023). We sought a ubiquitous, specific and non-intercalating marker for mtDNA in plant cells. The DNA binding protein TFAM has been used for this in human and yeast cells, however plant cells lack it and overexpression can impact mtDNA function (Alam et al., 2003; Kukat et al., 2011). Other DNA-binding proteins in plants are repair proteins, found in very low numbers per mitochondrion (Fuchs et al., 2020). Nucleoid organising proteins in plant mitochondria can also have preferential binding sites across the genome (Kim et al., 2021). We therefore were inspired by the work of (Deng et al., 2023), implementing the bacterial H-NS nucleoid binding protein, to bind each and every nucleoid in mitochondria. We confirmed that this nucleoid binding protein does not impact growth or respiration in Arabidopsis, and used it to validate the results seen in the SYBR stained cells. The photoconvertible nature of the Kaede protein can be used in future experiments to track the movement and colocalisation of mtDNA nucleoids originating in different individual organelles. This novel fluorescent protein construct will be particularly useful for imaging meaningful exchange of mtDNA nucleoids across all tissue types, elucidating further the role of mitochondrial dynamics on DNA maintenance and exchange.

We have shown that the complex physical and network behaviours of plant mitochondria are linked to whether the mitochondrion contains DNA or not. Mitochondrial motion is very costly in terms of the cell’s energy budget (each myosin step is estimated to use about 1 ATP for every 35nm travelled (Tominaga et al., 2003), and they can move at speeds of up to 10µm/s (Zheng et al., 2009)), suggesting the importance of this dynamic motion for cellular function. Much of the motion may be related to acquiring substrates and delivering products to different parts of the cell, and to and from other bioenergetic organelles. Individual mtDNA content is another parameter that affects their behaviour. The persistence of empty mitochondria and the factors connecting the presence of DNA to the outer membrane where movement is mediated are important remaining mysteries.

## Methods

### Data and Code availability

-Videos 1-4 are available at github.com/jo-c-bio/physical-genetic-dynamics -All code generated for this paper is also available at the link above, as are the scripts used to generate figures for this paper. Also available are .avi files of the two live time lapse collections for SYBR stained and mt-HI-NESS cells. Final plasmid sequences of AOX-mt-HI-NESS, IVD-mt-HI-NESS and Cox8-mt-HI-NESS are also available at the link above.

### Video Analysis

Video analysis closely follows (Chustecki *et al*., 2021). Time-lapse videos of SYBR stained mito-mCherry hypocotyl cells of 5-days old *Arabidopsis thaliana* were cropped to single cells using extracellular SYBR signal. 8 videos were taken, from which 10 unique cells were analysed (n=10). Ensuring resolution (4.83 pixels/µm) and time steps (2.21 secs) were consistent across the videos analysed, Trackmate v7.11.1 (Tinevez et al., 2017) in ImageJ v2.14.0 was used to track movement in the mito-mCherry channel only. Typical settings for SYBR stained cells were using the LoG Detector filter with blob diameter of 1.1µm, a threshold of 76-100, and a quality filter on spots if necessary of between 87-180. Then, the Simple LAP Tracker was applied with a linking max distance of 2.5, gap-closing distance of 3.5, and 2 frames used for gap-closing.

The same process was undertaken for live mito-mCherry AOX-mtHINESS cells, for 11 cells from 5 different seedlings (n=11). Again, resolution (3.8 pixels/µm) and time steps (2.52 secs) were consistent across the videos analysed. Tracking used slightly different parameters of blob diameter of 0.8µm, a threshold of 9.8-29.8 and a quality filter on spots if necessary of between 16.5-28.8, and the LAP Tracker linking max distance of 2.5, gap-closing distance of 3.5, and 1 frames used for gap-closing. For both video collections, manual editing of tracks was undertaken if deemed necessary.

Full .xml files were exported, and imported into RStudio (v2024.04.0+735) using the TrackmateR package (v0.3.10) (Royle, 2024), to include all spot statistics. To define trajectories as having or not having mtDNA, we use the second channel of the SYBR/mt-HI-NESS signal, and two definitions-Contrast and Maxima. Contrast is calculated as the Intensity (A.U) of pixels inside the spot radius (*r*) compared to outside as defined by 2*r* by (I_in_-I_out_)/(I_in_+I_out_), and for each cell, the contrast cutoff for what counted as a candidate SYBR/mt-HI-NESS signal was manually selected using the visualisation tool and spot pseudo-colouring in Trackmate. Cutoffs ranged from 0.05-0.22 contrast values, where anything below was without mtDNA, and anything above was with mtDNA.

For Maxima definition of mtDNA presence, the FindMaxima (Wagner, 2016) function in ImageJ was run through a macro, across three frames (1,50,100) of each video, and looped over prominence values in increments of 50 from 150-500, with the ‘strict’ setting applied. Final prominence values used across the videos ranged from 200-400 for SYBR videos, and 10 or 11 for mt-HI-NESS videos. Images with maximum points as crosses were viewed to choose representative best prominence value for true SYBR/mt-HI-NESS signal, and this chosen prominence value was used for each video to generate a list of pixel coordinates from puncta that were SYBR/mt-HI-NESS signal. After conversion to microns to match Trackmate output, the R point.in.polygon function from sp package v2.1-4 was used to assess whether or not the maxima puncta sat inside a Trackmate spot from the mito-mCherry signal. If it did, the mitochondrion at that frame has mtDNA according to our Maxima definition, and if not, it was without mtDNA.

For both Maxima and Contrast definitions, three strictness definitions were then defined. Each video had its own contrast and prominence value. We define the first as *Lenient* in that only one frame in the entire trajectory needs to be above the threshold value to pass (as this mitochondrion having mtDNA). The next is *tolerant*, where at least two adjacent frames must be above the threshold to pass. The last definition is *strict*, where at least three adjacent frames must be above the threshold to pass.

Physical statistical analysis includes speed (µm/s), taken as distance moved per frame per trajectory. Area (µm^2^) is taken as the region inside a polygon plotted using the furthest points of a trajectory of frame length ⩾3 (convex hull area). Minimum intermitochondrial distance (µm) is the minimum Euclidean distance between each mitochondrion, and any other mitochondrion in the cell, per frame. Any log values represent values at log base *e*.

We characterise encounter networks as undirected edges between nodes representing mitochondria. Edges are built when mitochondria are within 1.6µm of each other, and do not represent fusion events. Encounter networks build as frame times progress, and network build-up is historical. Node degree is then calculated as the number of immediate neighbours each node is connected to. This analysis aimed to investigate connectivity values for nodes with and without a characteristic, therefore we did not implement global values of connectivity for entire networks, as we analysed connectivity between individual nodes, instead of between whole networks, and avoided splitting the network into subgraphs. R packages iGraph (v2.0.3) and brainGraph (v3.1.0) were used to calculate network statistics and visualise networks.

### Plant growth

Seeds of Col-0, mito-mCherry, mtGFP, mito-mCherry mt-HI-NESS and Col0 mt-HI-NESS lines were surface sterilised using 50% (v/v) household bleach solution with continual inversion before washing three times with sterile ddH_2_O, and plating onto ½ MS agar with 5µg/mL Nystatin at pH 5.7 before stratification at 4°C for two days in the dark. Seedlings were either grown to 4-5 days old in ½ MS before use or transplanted to Berger BM2 germinating soil with Turface MVP. Any plants grown on soil were grown at 22°C in 8 hrs dark,16 hrs light.

*N. benthiamiana* plants were grown in greenhouse conditions with supplemented light 12hrs a day and temperature ranges at 27-24°C daytime and 21-19°C nighttime from seed until ∼3-4 weeks old.

### Plant tissue fixation staining and imaging

5-days old Arabidopsis seedlings were fixed in 3.5% paraformaldehyde (EMS, #15710) in PBS buffer (Gibco, #10010023) for 30 mins, ensuring complete submersion by applying gentle vacuum pressure to the solution. Seedlings were washed with PBS before further staining.

SYBR green (invitrogen, #S7563) was used at a 1:1000 dilution in ddH_2_O, and 5-days old Arabidopsis seedlings were stained for 45 mins before washing in PBS or H_2_O and use.

All laser scanning confocal micrographs with the exception of the 14 mito-mCherry AOX-mtHINESS time lapse videos were taken on an upright Nikon A1-NiE system (RRID:SCR_020318) using NIS-Elements Confocal image acquisition and analysis software, using a 60x water objective. Images were taken using a sequential channel series setting, to minimise cross-channel signal, and the channels used were: GFP, Ex: 488nm, Em: 525 (500-550), TxRed, Ex 561, Em: 595 (570-620), Cy5, Ex: 640nm, Em: 700 (663-738).

All time-series images taken on the Nikon system have a frame time of 2.21s.

For imaging of live hypocotyl cells, and fixed hypocotyl and leaf epidermal cells, 5-days old seedlings grown on plates were used. For mt-HI-NESS leaf epidermal images, ∼2/3 week-old plants were used. Tissues were mounted in water on glass slides, with double sided sticky tape and a glass cover slip.

For photoconversion of the Kaede fluorescent protein of the AOX/IVD-mt-HI-NESS lines, the A1 and ND Stimulation features of the NIS-Elements were used to target 4% of the total field of view for 5 seconds at 10% of 20mW LD 405nm laser power. Frame intervals were 2.15secs and the total time imaged was 95 secs.

The 14 mito-mCherry AOX-mtHINESS time lapse videos analysed were of single hypocotyl cells of 5-day of seedlings, taken on an inverted Zeiss LSM 980 with ZEN blue image acquisition and analysis software, using a 63x oil immersion lens. Time lapses were taken in two tracks, GFP, Ex: 488nm, Em: 509 nm (490-561nm), mCherry, also capturing chloroplast autoflourescence, Ex: 587, Em: 610nm (596-685nm). This collection of time series has a frame time of 2.52 secs.

### Cell segmentation with IMARIS

Z-stack files of fixed cells were imported into IMARIS (version 10.1.0). The cell feature was used to define thresholds for segmenting vesicles (mitochondrial DNA puncta) and nuclei (mitochondria). The cell outline segmentation was not implemented, as a reliable cell wall/membrane stain could not be implemented. To work around this, manual cropping of cell outlines based on SYBR stain at the cell wall was performed in ImageJ2 (2.14.0/1.54f) before import. Further cropping was done using cropping planes in IMARIS, until a clear view of a single cell in each case was achieved. In most cases, the same thresholding parameters were used, briefly: Nucleus Smooth Filter Width = 0.2µm, Nucleus Background Subtraction Width = 0.25-0.4 µm, Nucleus Manual Threshold Range: 77.3-261.3, Vesicle Estimated Diameter = 0.600 µm, Vesicle “Quality” above range: 63.5-207. The hypocotyl hyp1 sample also had an extra filter of “Nucleus Number of Voxels” above 2.05.

During analysis of exported data, any segmented ‘mitochondrion’ < 0.1µm^3^ was removed. SYBR signals from the nucleus and chloroplasts were avoided by quantifying only mtDNA puncta within segmented mitochondria.

### Correlation Analysis

Spearman correlation was calculated using the cor.test function of the stats (version 3.6.2) R package, using Spearman’s *rho* statistic, for a rank-based association between values, and which uses p-values computed using algorithm AS 89, set out in (Best and Roberts, 1975).

### mt-HI-NESS plasmid construction

The mt-Kaede-HI-NESS construct was kindly provided by Professor Timothy Shutt, University of Calgary, developed for use in U2OS human cells, using cox8 as the MTS (mitochondrial targeting sequence). This construct was transformed into TakaraBio Stellar *E. coli* cells (TakaraBio, #636763) and grown on LBA plates with Ampicillin (100µg/mL).

Plasmid DNA was extracted using a column-free plasmid prep method (Green & Sambrook, 2012), and mt-Kaede-HI-NESS-HA was amplified from the plasmid (linearised using AvrII restriction enzyme (NEB, # R0174S) using RP (5’-CAAGAAAGCTGGGTTTAAGCG-3’) and FP2 primers (5’-CACCGCCACCATGTCCGT-3’) resulting in only one cox8 sequence being carried through, and a CACC sequence for directional TOPO cloning being introduced. This 1003bp PCR product was extracted from a 1% agarose gel using the TakaraBio gel extraction kit (TakaraBio, #740609.50), and introduced to the pENTR TOPO cloning vector (Kan^R^) (Invitrogen, #K240020) which contains the attL1 and attL2 sites LR clonase reactions, and transformed into OneShot Top10 competent *E.coli* (Invitrogen, #C404003).

In parallel, the cox8 MTS was switched for the plant-specific AOX or IVD MTS, and AOX/IVD-mt-Kaede-HINESS-HA constructs were synthesised by ThermoFisher GeneArt (https://www.thermofisher.com/us/en/home/life-science/cloning/gene-synthesis/geneart-gene-synthesis.html), following codon optimisation for Arabidopsis using the VectorBuilder Optimisation tool (https://en.vectorbuilder.com/tool/codon-optimization.html), and with attL1 and attL2 sites introduced for the LR clonase reaction into *Agrobacterium tumefaciens*. These were shipped in pMA-RQ plasmids (Amp^R^), and were transformed into TakaraBio Stellar *E. coli* cells, before plasmid DNA extraction.

The three plasmid types, cox8/AOX/IVD-mt-Kaede-HINESS-HA were inserted into separate pUB-DEST vectors via the LR clonase reaction (Spec^R^) (Invitrogen, #11791020), transformed into Top10 competent *E.coli,* sequenced and verified. This puts the inserted construct under control of the UBQ10 promoter. After one final plasmid prep, they were all transformed into *A. tumefaciens* strain C58C1 for use in infiltration and floral dipping. At each step, plasmids were sequenced using Eurofins Genomics whole plasmid sequencing service.

### Transient infiltration of *N. benthiamiana*

C58C1 *A. tumefaciens* lines were grown in YEBS + RGS for 2 days, alongside a mitochondrial-targeted red fluorescent protein IVD-FP611 (in pZP212). Following the protocol previously described (Mangano, Gonzalez and Petruccelli, 2014), ∼4 week old *N. benthiamiana* leaves were syringe infiltrated on the abaxial surface with bacterial preparations (OD = 0.35 of each line). Plants were left to recover in a growth chamber for 48 hours before imaging.

### Stable transformation of *A. thaliana*

∼4 week old Col-0 and mito-mCherry plants were transformed via floral dip with *A. tumefacien* carrying either AOX or IVD-mt-Kaede-HINESS-HA constructs. A previously published protocol was followed (Zhang *et al*., 2006), using a 5% (wt/vol) sucrose solution at pH 5.6 to resuspend cultured bacteria, and a second dip was performed 7 days after the first. Seeds were left to develop before screening with BASTA on soil.

### BASTA® treatment for transformed Arabidopsis screening

Glufosinate Ammonium (BASTA®, plantmedia.com) was diluted to 300µM in a spray bottle with ddH_2_O and sprayed upon seeds before stratification, then also to newly emerged seedlings and 1-2 week old plants, removed from growth chambers. Col-0 (susceptible) and pUBdest transformed (resistant) plants were grown alongside to monitor the effectiveness of treatment. Once sprayed, trays were left with plastic domed lids on for 8 hours before returning to the growth chamber.

### Root and rosette assays

Seeds of each genotype were bleach sterilised and sown onto square ½ MS plates, after stratification at 4°C for 2 days, placed at a 70° angle in a growth cabinet and grown for 9 days without disturbance. Images were taken using the brightfield setting on a GelDoc Imager. For Col-0 background root comparison, the same method was used but seedlings were grown on 1/2 MS with sucrose and imaged using a Nikon camera. For all, root lengths were hand traced from cotyledon base to root tip with the flexible line tool in ImageJ, after setting the scale based on the plate size, lengths were exported and further analysed in R.

For rosettes, seeds were grown on upright plates for 11 days before sowing on soil and growing for another 19 days. Rosette images were traced using freehand draw in ImageJ and the area was taken after setting the scale for pot size, exported and analysed in R.

### Immunodetection of mitochondrial proteins

For each line, 50mg of 23-day-old *Arabidopsis* seedlings were frozen in dry ice (kindly provided by Jithesh Vijayan), ground in 2mL cold homogenisation buffer (40mM HEPES, 0.4M D-sorbitol, 2mM EDTA, 50µM PVP40, 14.4mM ß-mercaptoethanol), passed once through Miracloth, and once through a 100µm filter to extract the total homogenate, centrifuged at 17,200g for 30mins at 4°C, resuspending each in 30µL 1X Laemmli Buffer (BioRad #1610747).

Standard immunodetection techniques were followed using 10µL of each sample loaded into miniprotean TGX stain-free gels (BioRad #4568094), using dilute loading buffer (BioRad #1610732), and Precision Plus marker (10-250kDa) (BioRad #1610374), and run at 200V until separation, and imaged via GelDoc Go.

Following migration, proteins were transferred onto a nitrocellulose membrane (BioRad, #1620213), using the BioRad trans-blot (#1704150), blocked in EveryBlot Buffer (BioRad, #12010020) and incubated in primary antibodies from Agrisera AB (Sweden): CoxII at 1:1000 (#AS04053A), AtpB at 1:1000 (#AS163976), PsbA at 1:10,000 (#AS05084) and H3 at 1:5000 (#AS10710), then goat anti-rabbit secondary antibody from Southern Biotech (USA) (#4050-04). Final color detection was via the AP conjugate substrate kit (BioRad #1706432), and imaged via GelDoc Go.

### Oxygen consumption measurements

∼60 (<u>±</u>1) mg of leaf discs (7mm width) were cut from the middle tip region of 25-day-old *Arabidopsis* leaves and placed in respiratory activity incubation medium (10mM HEPES, 10mM MES, 2mM CaCl_2_, pH 7.2) for 30 mins in the dark. Oxygen consumption measurements were taken using the Oxygraph+ oxygen electrode system (Hansatech Instruments, Pentney, Norfolk, UK), following (Sew et al., 2013). Calibration by removal of all oxygen in 1mL ddH_2_O was performed using an excess of Sodium dithionate. Oxygen concentration was read over 15mins in the dark in 2mL of respiratory activity incubation medium, and oxygen consumption rate was calculated as nmol O_2_ per minute per gram of fresh weight (FW).

## Acknowledgements

We thank Prof. You (Joe) Zhou and Dr. Bara Altartouri of the Microscopy Core at the Nebraska Centre for Biotechnology for their invaluable help and microscopy advice. Thank you to Dr. Heather Jensen-Smith at the University of Nebraska Medical Centre Advanced Microscopy Core for access and advice on the IMARIS software. Thanks to Prof. Timothy Shutt (University of Calgary) and his team for the kind gift of the mt-HI-NESS construct. We thank Dr. Huang Li and Prof. Jeff Mower (University of Nebraska-Lincoln) for the use of the Hansatech Oxygraph+ system. We thank Prof. Christian Elowsky for his imaging advice and encouragement, and Dr. Kostas Giannakis for fruitful conversations on network analysis. This work was funded in part by a grant from the National Science Foundation to ACC (MCB-1933590). Major support was from a University of Nebraska Foundation fund in memory of Frank and Edith Christensen.

## Author contributions

JMC and ACC designed the research. AQS and MHF conducted experiments. JMC conducted experiments, performed imaging, image analysis and data analysis. JMC and ACC wrote the manuscript. All authors edited the manuscript.

## Supplementary material

**Video 1:** Time lapse video of SYBR stained hypocotyl cell of 5-day old Arabidopsis seedling. Frame interval = 2.21 secs, Exported at 10 fps. Green channel-SYBR signal (mtDNA). Red Channel - mito-mCherry signal (mitochondria). https://github.com/jo-c-bio/physical-genetic-dynamics/blob/main/SupplementaryVideos/Video1-SYBRhypocotyl.avi

**Video 2:** mito-mCherry IVD-mt-HI-NESS timelapse. Time-lapse video of leaf epidermal cells of 2-3 week old mito-mCherry Arabidopsis stably transformed with IVD-mt-HI-NESS, Frame interval = 3.28 secs, exported at 10 fps. Green; IVD-mt-HI-NESS signal (mtDNA), Red Channel; mito-mCherry signal (mitochondria), Blue; Chloroplast autofluorescence, https://github.com/jo-c-bio/physical-genetic-dynamics/blob/main/SupplementaryVideos/Video2-mCherry-IVD-mtHINESS.avi

**Video 3:** mito-mCherry AOX-mt-HI-NESS timelapse. Time-lapse video of leaf epidermal cells of 2-3 week old mito-mCherry Arabidopsis stably transformed with AOX-mt-HI-NESS, Frame interval is 6.06 secs, exported at 10 fps. Green; AOX-mt-HI-NESS signal (mtDNA), Red Channel; mito-mCherry signal (mitochondria), Blue; Chloroplast autofluorescence, https://github.com/jo-c-bio/physical-genetic-dynamics/blob/main/SupplementaryVideos/Video3-mCherry-AOX-mtHINESS.avi

**Video 4:** Col-0 IVD-mt-HI-NESS photoconversion time-lapse. Time-lapse of photoconversion from green to red of IVD-mt-HI-NESS (mtDNA marker) in Col-0 Arabidopsis epidermal leaf around 2-3 weeks old with no mitochondrial marker. The first frame shows the region hit by a 405nm laser, frame interval is 2.15 secs, and there is a 5 secs gap between frames 3 and 4 when the laser was applied. Total time elapsed 95 secs. Exported at 2 fps. https://github.com/jo-c-bio/physical-genetic-dynamics/blob/main/SupplementaryVideos/Video4-IVD-photoconversion.avi

---

**Figure S1:**
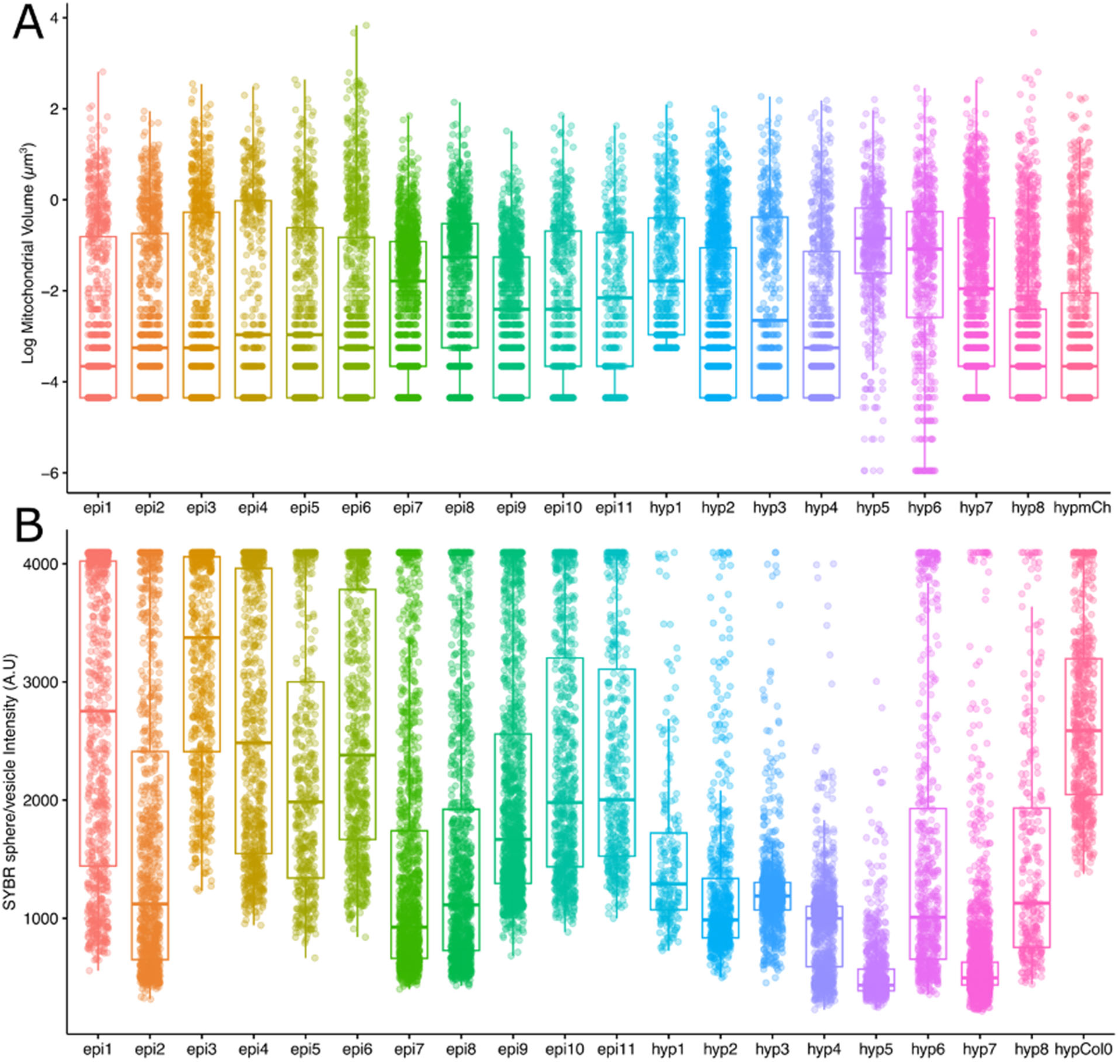
Statistics from fixed, SYBR stained leaf epidermis n = 11, hypocotyl = 8 cells and controls. A) Log mitochondrial volume (µm^3^) extracted from fixed mito-mCherry cells compared to n = 1 hypocotyl from an unstained mito-mCherry seedling. B) Intensity of segmented SYBR spheres (representing the mean intensity of all voxels in the sphere) from fixed mito-mCherry cells, compared to n = 1 stained Col-0 hypocotyl (no mitochondrial marker).

**Figure S2:**
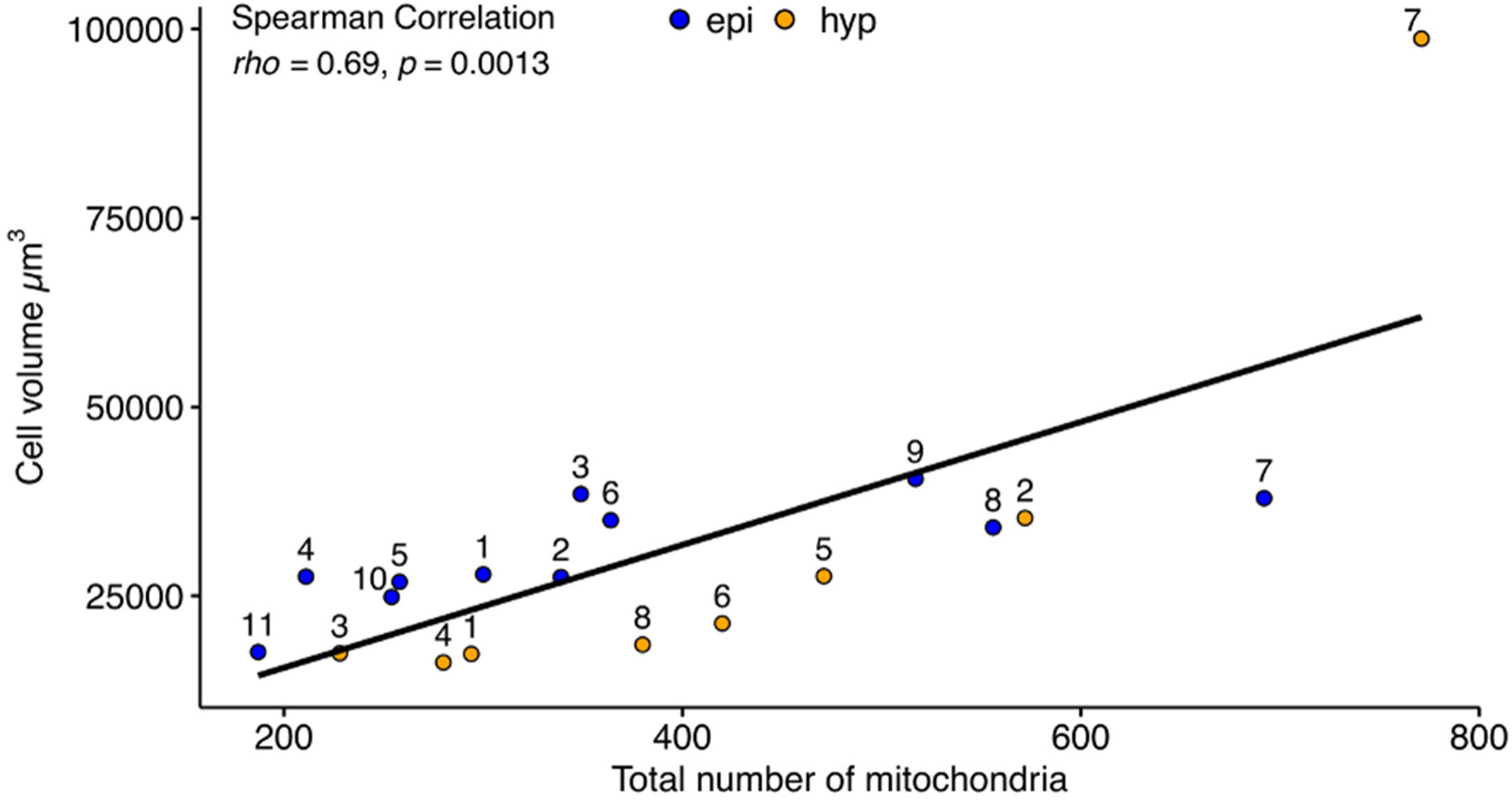
Spearman correlation between cell volume and total mitochondrial number, across the two cell types used (leaf epidermis n = 11, hypocotyl n = 8), with a linear regression line added.

**Figure S3:**
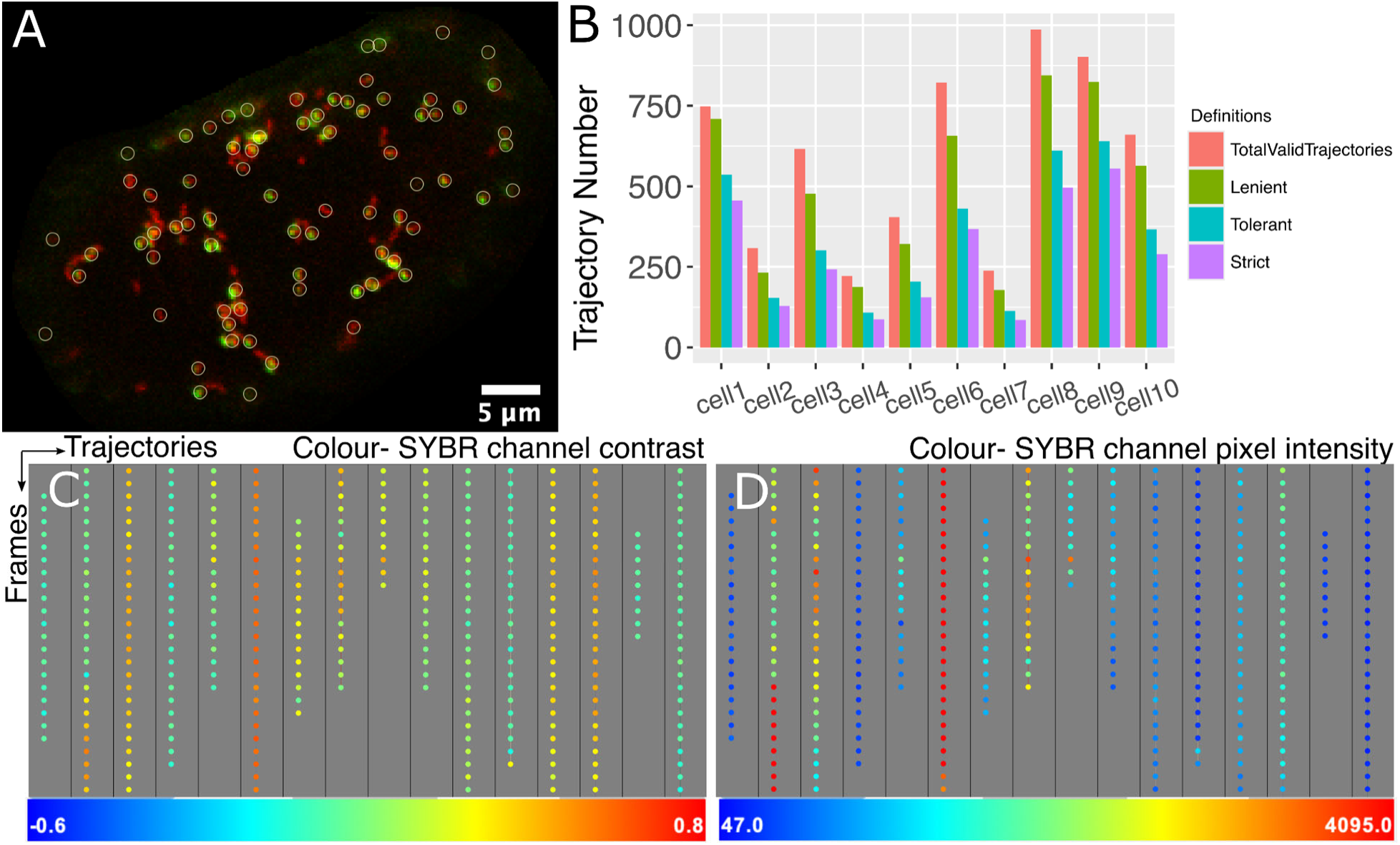
Detecting and defining mtDNA presence using contrast. A) Single Arabidopsis hypocotyl cell with mito-mCherry marker, stained with SYBR green, still frame from time-lapse (same as in Figure 2A), with detected mitochondria (red) using Trackmate (white circles, diameter 1.1µm), showing only those that passed the chosen Contrast threshold for this time-lapse. B) Quantification of trajectory number over total, alongside three definitions of mtDNA presence, in order of strictness, across 10 single hypocotyl cells. C,D) Trackschemes, from Trackmate (Tinevez *et al*., 2017) showing multiple trajectories (across), made up of points at each time frame (down), colour coded by contrast value of SYBR channel (C), or pixel intensity values of SYBR channel (D). Lower values = blue, higher values = red.

**Figure S4:**
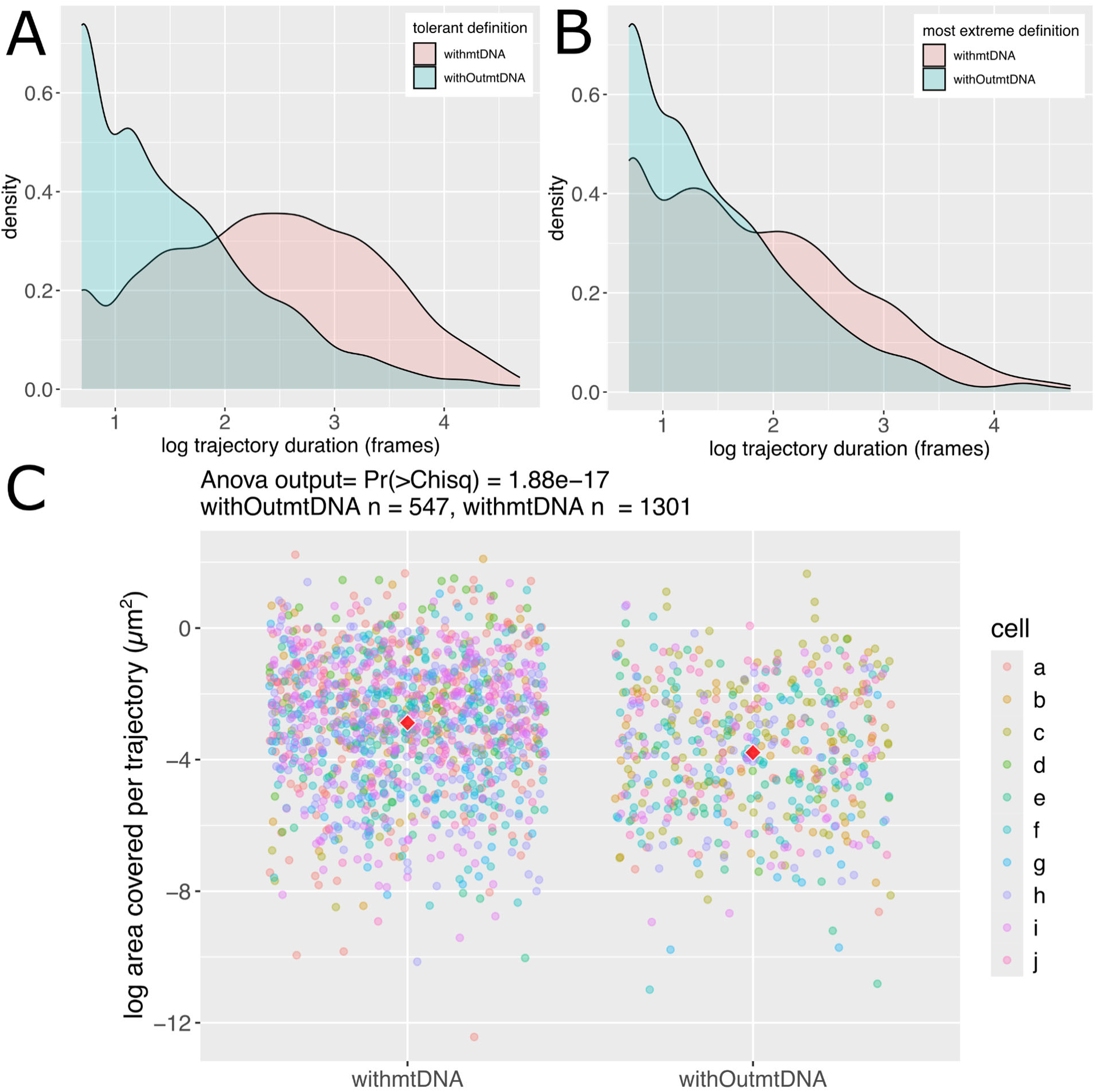
Impact of differing mtDNA presence definitions on area analysis. Comparing trajectory duration in frames between trajectories defined as without (blue) or with (pink) mtDNA according to a ‘tolerant’ (A) or ‘strictest’ (B) definition. C) Impact of *most extreme* implemented definition of mtDNA presence on differences in cell area travelled by trajectories (µm^2^). P-value is the outcome of a blocking factor Anova (cells as block). Red diamonds represent the mean of all points.

**Figure S5:**
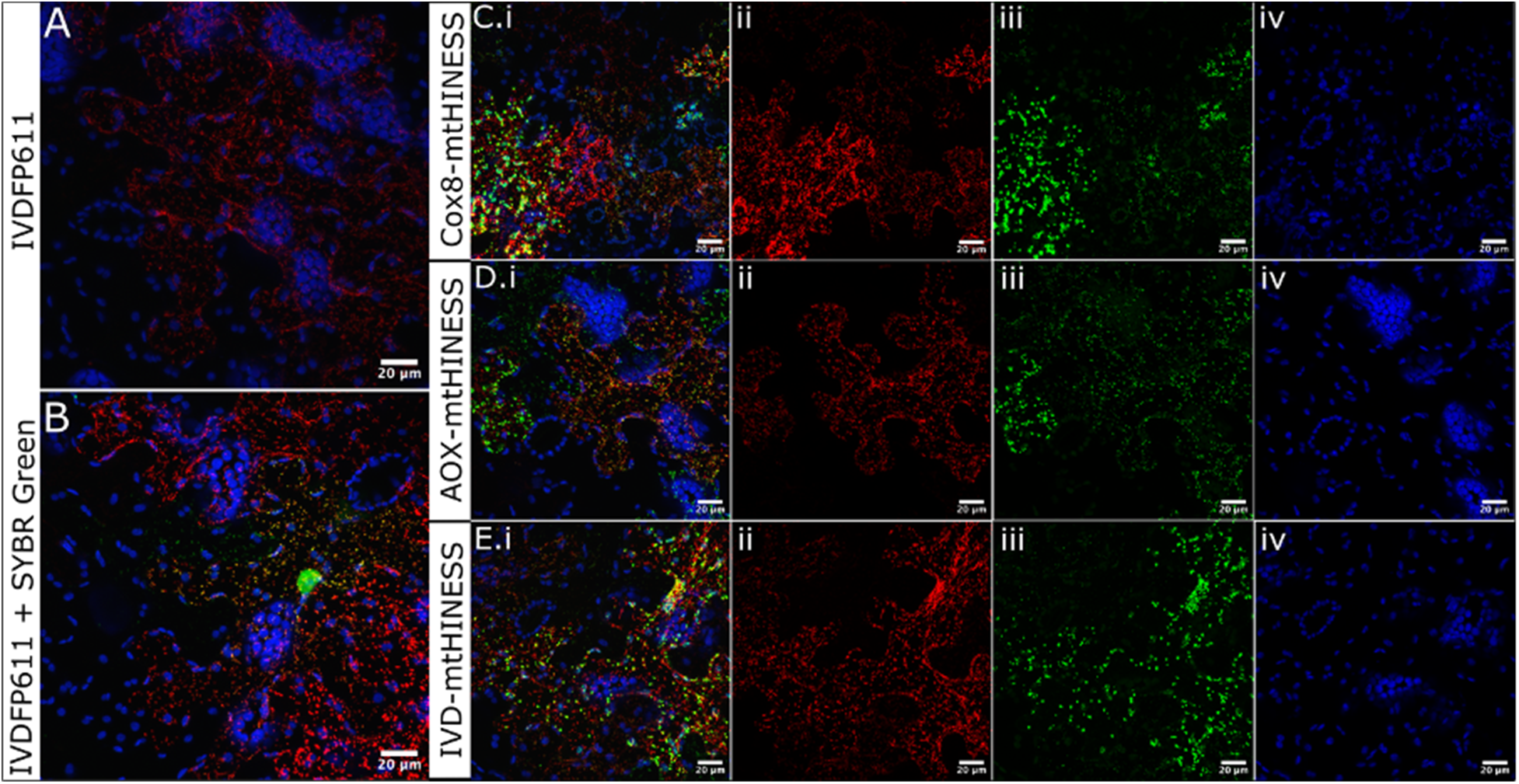
Validation of mt-HI-NESS nucleoid reporter constructs in *N. benthamiana* transient infiltrations, abaxial epidermal cells. A) Expression of IVD-FP611 control for mitochondrial signal (red) and chloroplast autofluorescence (blue). B) Expression of IVDFP611 mitochondrial localised fluorescent protein (red) subsequently stained with SYBR green (green), with chloroplast autofluorescence (blue). C-E) Merged (i) coinfiltrations of IVDFP611 (ii, red) with mt-HI-NESS constructs (iii, green) differing by mitochondrial targeting sequence Cox8 (C), AOX (D), IVD (E) respectively, with chloroplast autofluorescence also shown (iv, blue).

**Figure S6:**
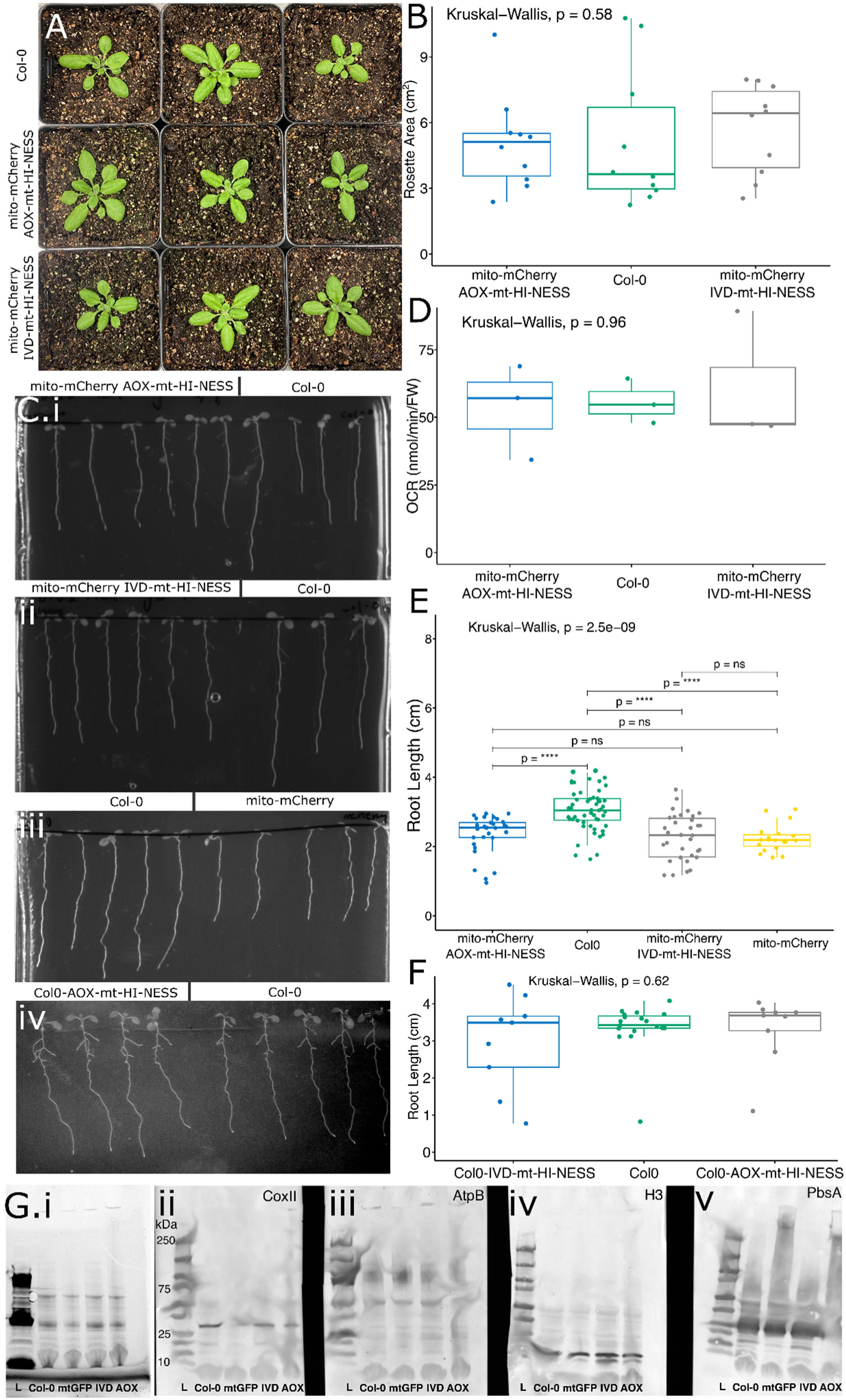
Physiological validation of mt-HI-NESS Arabidopsis stable transformants. A) Rosette images of Col-0 (WT) and mito-mCherry background transformants, n=3 B) Rosette area analysis between the same genotypes as in (A), where n=10 for each line. C) Root growth assays comparing (i) mito-mCherry AOX-mt-HI-NESS and Col-0, (ii) mito-mCherry IVD-mt-HI-NESS and Col-0, (iii) Col-0 and mito-mCherry, and (iv) Col-0 AOX-mt-HI-NESS and Col-0. D) Oxygen consumption rate (OCR) in (nmol/min/FW) across Col-0 WT, mito-mCherry IVD-mt-HI-NESS and mito-mCherry AOX-mt-HI-NESS. E) Quantification of root length (cm) across the four lines seen in (C), mito-mCherry AOX-mt-HI-NESS n=32, Col-0 n= 54, mito-mCherry IVD-mt-HI-NESS n= 34, mito-mCherry n=19 F) Quantification of root length (cm) in the Col-0 background, Col-0 AOX-mt-HI-NESS n= 9, Col-0 n=19, Col-0 IVD-mt-HI-NESS n= 9. G) Immunoblots of Col-0, mtGFP, mito-mCherry IVD-mt-HI-NESS (IVD), and mito-mCherry AOX-mt-HI-NESS (AOX) detecting mitochondrial, nuclear and chloroplast proteins showing gel loading control (i), CoxII (mitochondrial, ii), AtpB (mitochondria, iii), H3 (nuclear, iv), PsbA (chloroplast, v), L = ladder (10-250kDa). *P*-values represent Kruskal–Wallis test outcomes across all genotypes, and pairwise *P*-values are false discovery rate-adjusted outcomes of a post-hoc Dunn test, ****= p< 0.001, ns = not significant. Boxplots represent the median and 25th/75th percentile, with whiskers showing the smallest/largest value within 1.5× the interquartile range.

**Figure S7:**
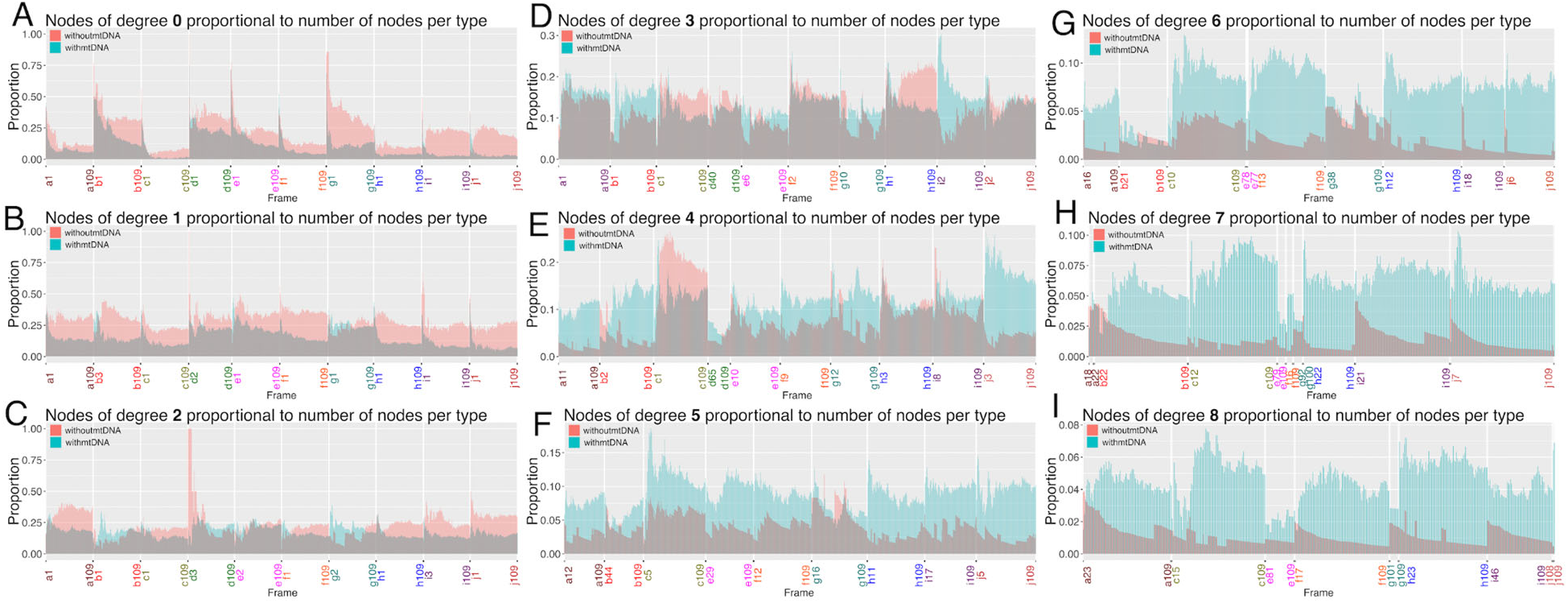
Number of nodes with degree value 0-8 (A-I), proportional to the number of nodes of equal mtDNA status, plotted over n = 10 SYBR stained hypocotyl cells, color coded on the x-axis, and all networks from all frames. Not all networks at individual frames had high degree values, so for graphs of higher degree, not all frame times or cells will be represented. First and last viable frames for each time-series noted on x-axis.

**Figure S8:**
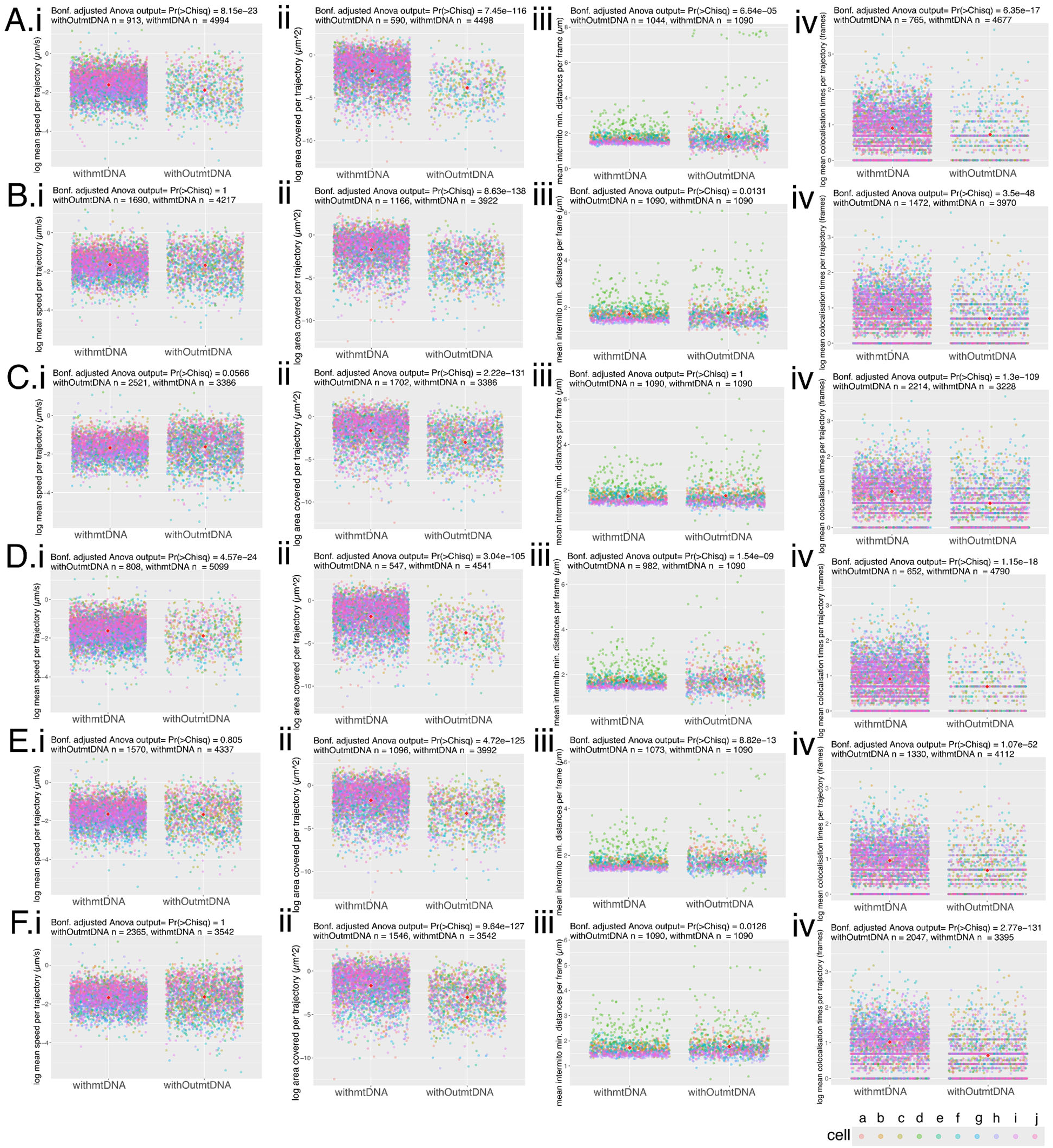
Physical statistics reported for SYBR stained mito-mCherry hypocotyl cells, across all mtDNA foci definitions and all presence/absence trajectory definitions. Foci were defined by either Contrast (A-C) or Maxima (D-F) of the SYBR signal, and the presence/absence of mtDNA in each trajectory was defined by lenient (A,D), tolerant (B,E) or strict (C,F). Statistics shown are those described in the main text i) log mean speed per trajectory (µm/s), ii) log area per trajectory (µm^2^), iii) mean intermitochondrial distance per trajectory (µm), (iv) log mean colocalisation time per trajectory (frames). n= 10 cells, each colour coded with key at lower right. P-values are the outcome of a blocking factor Anova (cells as block), adjusted across all statistics tested using the Bonferroni method. Red diamonds represent the mean of all points.

**Figure S9:**
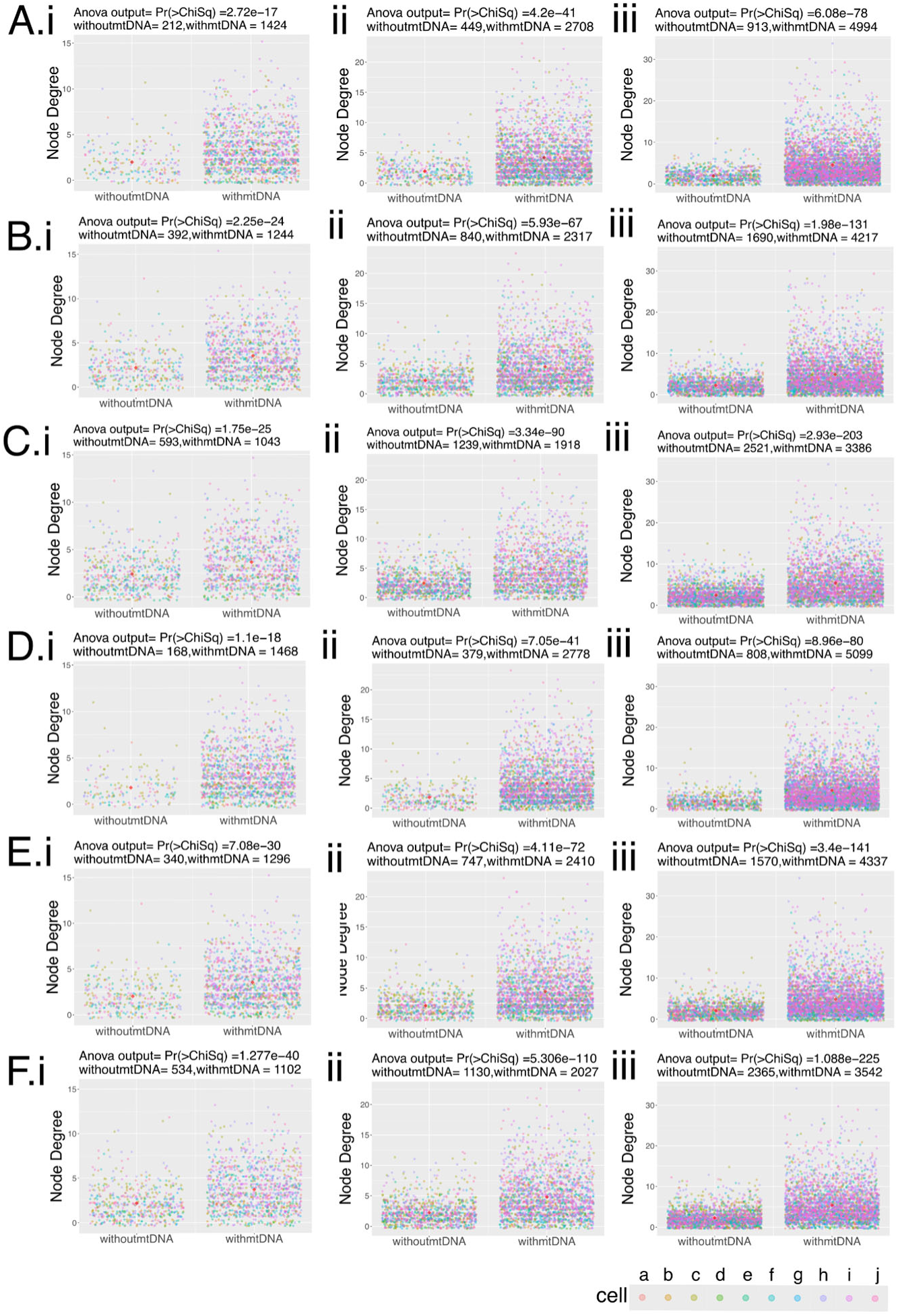
Degree connectivity statistics reported for SYBR stained mito-mCherry hypocotyl cells, across all mtDNA foci definitions and all presence/absence trajectory definitions. Foci were defined by either Contrast (A-C) or Maxima (D-F) of the SYBR signal, and the presence/absence of mtDNA in each trajectory was defined by lenient (A,D), tolerant (B,E) or strict (C,F). Degree, as defined in text, the number of immediate neighbours of each node, here split by presence/absence of mtDNA, and shown for networks at frame 20 (i), 50 (ii) and 109 (iii). n= 10 cells, each colour coded with key at lower right. P-values are the outcome of a two-way blocking factors anova (cells as block). Red diamonds represent the mean of all points.

**Figure S10:**
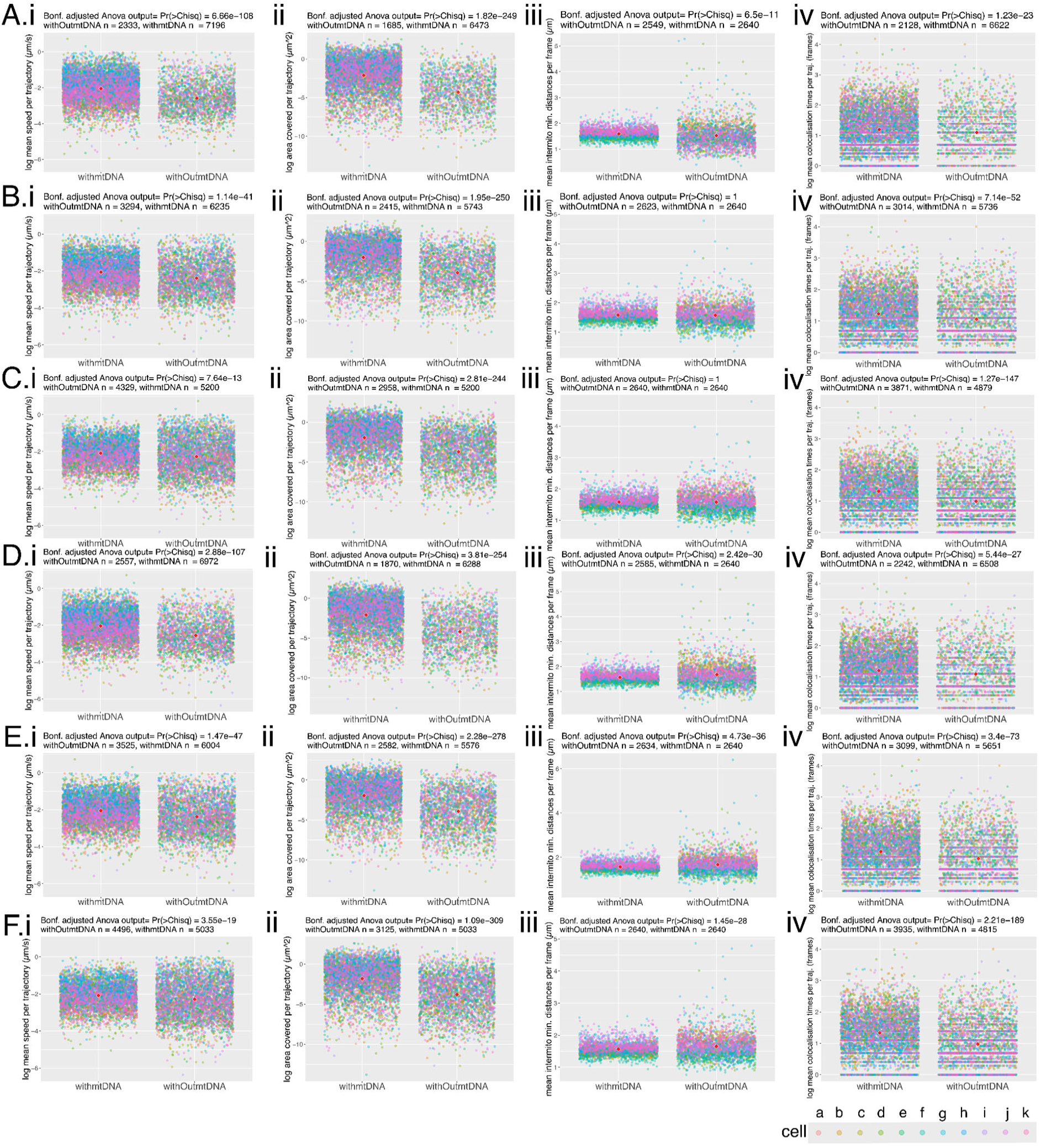
Physical statistics reported for mito-mCherry AOX-mt-HI-NESS hypocotyl cells, across all mtDNA foci definitions and all presence/absence trajectory definitions. Foci were defined by either Contrast (A-C) or Maxima (D-F) of the SYBR signal, and the presence/absence of mtDNA in each trajectory was defined by lenient (A,D), tolerant (B,E) or strict (C,F). Statistics shown are those described in the main text i) log mean speed per trajectory (µm/s), ii) log area per trajectory (µm^2^), iii) mean intermitochondrial distance per trajectory (µm), (iv) log mean colocalisation time per trajectory (frames). n= 11 cells, each colour coded with key at lower right. P-values are the outcome of a blocking factor Anova (cells as block), adjusted across all statistics tested using the Bonferroni method. Red diamonds represent the mean of all points.

**Figure S11:**
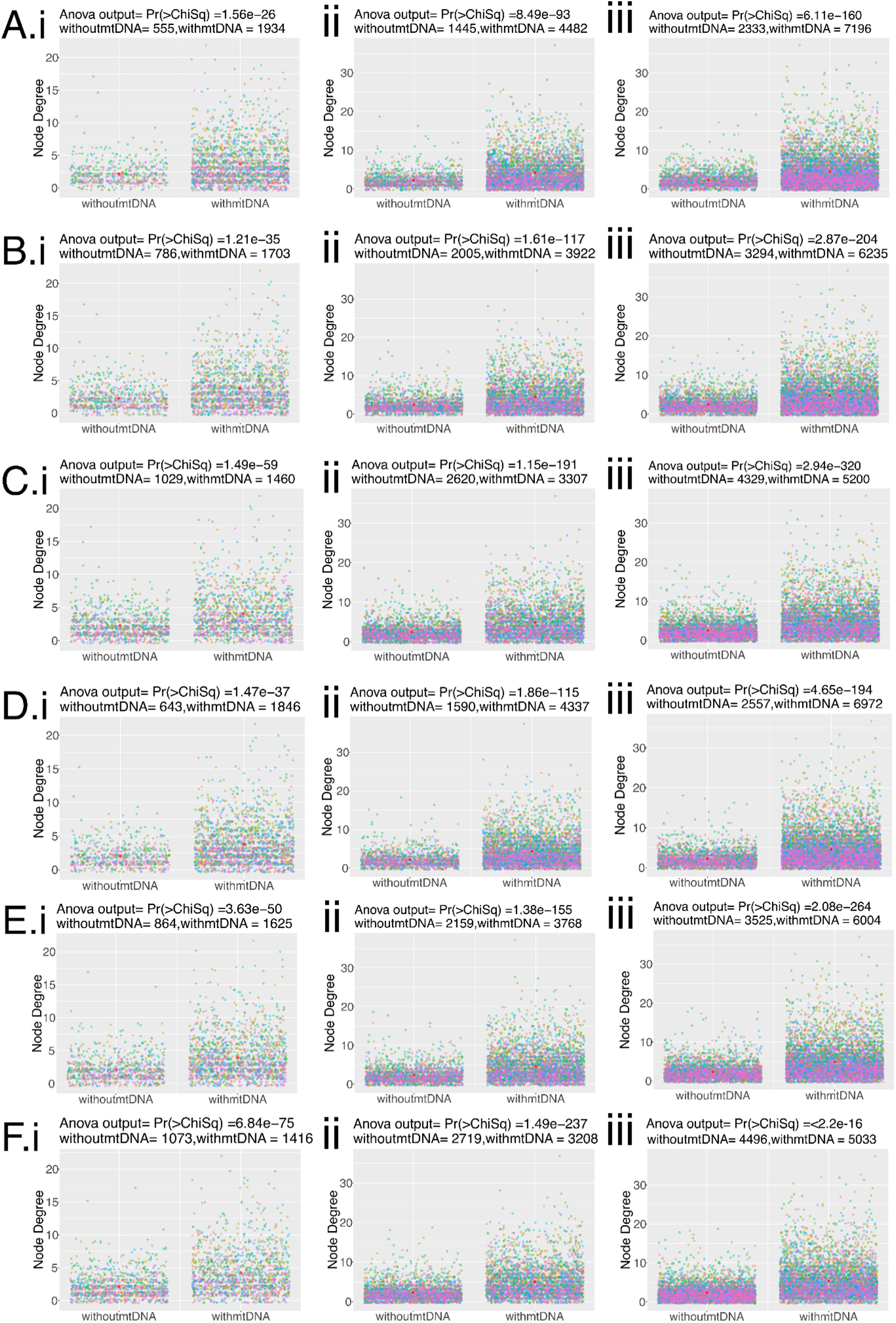
Degree connectivity statistics reported for mito-mCherry AOX-mt-HI-NESS hypocotyl cells, across all mtDNA foci definitions and all presence/absence trajectory definitions. Foci were defined by either Contrast (A-C) or Maxima (D-F) of the mt-HI-NESS nucleoid signal, and the presence/absence of mtDNA in each trajectory was defined by lenient (A,D), tolerant (B,E) or strict (C,F). Degree, as defined in text, the number of immediate neighbours of each node, here split by presence/absence of mtDNA, and shown for networks at frame 50 (i), 150 (ii) and 240 (iii). n= 11 cells, each colour coded with key at lower right. P-values are the outcome of a two-way blocking factors anova (cells as block). Red diamonds represent the mean of all points.

**Figure S12:**
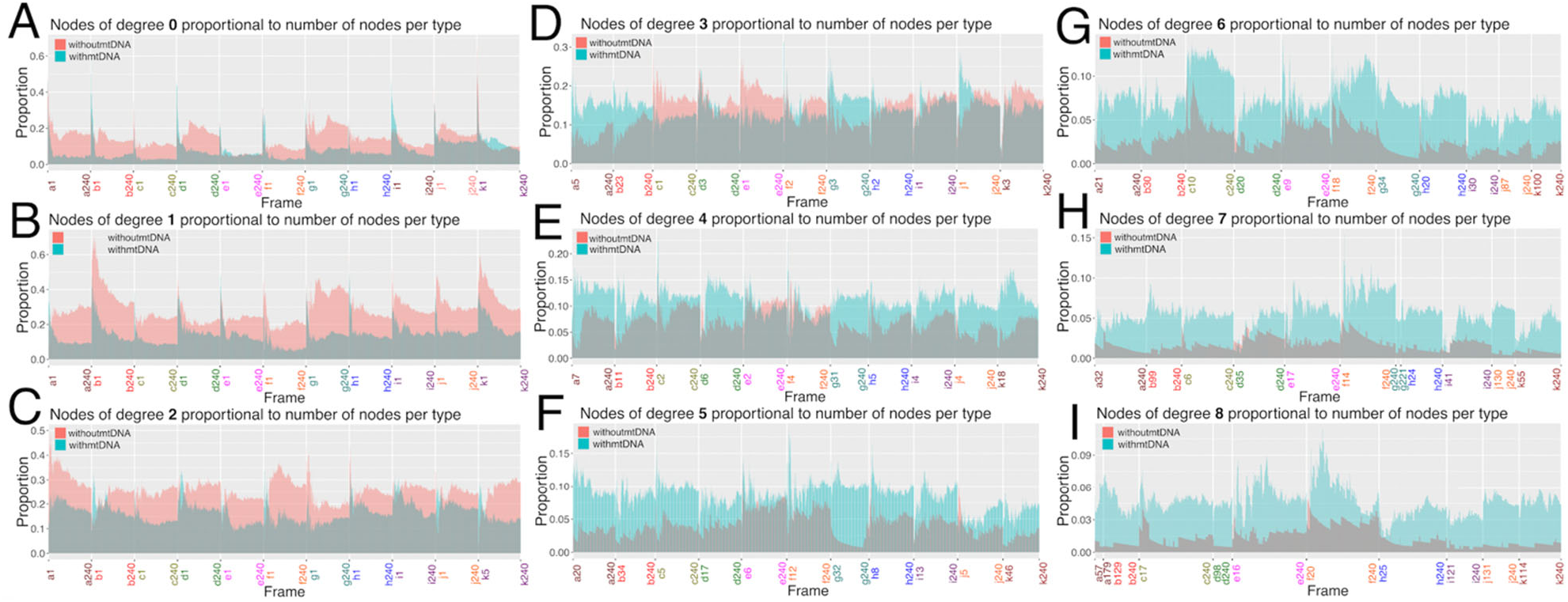
Number of nodes with degree value 0-8 (A-I), proportional to the number of nodes of equal mtDNA status, plotted over n = 11 mito-mCherry AOX-mt-HI-NESS hypocotyl cells, color coded on the x-axis, and all networks from all frames. Not all networks at individual frames had high degree values, so for graphs of higher degree, not all frame times or cells will be represented. First and last viable frames for each time-series noted on x-axis.

